# The economic strategies of superorganisms

**DOI:** 10.1101/2025.02.21.639603

**Authors:** Lily Leahy, Hannah L. Riskas, Ben Halliwell, Ian J. Wright, Amelia G. Carlesso, Nathan J. Sanders, Tom R. Bishop, Catherine L. Parr, Steven L. Chown, Jonathan Z. Shik, Alan N. Andersen, Heloise Gibb

## Abstract

The leaf economics spectrum links strategies of plant investment in resource-acquiring leaves to overall fitness. We test whether an economic spectrum can also explain variation in ecological strategies of ant species across environmental gradients, where colony investment in workers is analogous to plant investment in leaves. A fast return of resource investment was associated with large colonies of smaller, less robust, short-lived workers with low nitrogen:phosphorus ratios. Slow resource payback was associated with small colonies of densely built, energetically conservative and longer-lived workers with high nitrogen:phosphorus ratios. Species representing the entire economic continuum co-occurred in all communities. Phylogenetic analyses suggest genus level conservation of core investment templates. These results unify studies of plants and ants, suggesting common economic principles apply across the tree of life.

## Main Text

One of the basic principles common to all life is that organisms are sustained by the storage and turnover of energy for fueling metabolism and chemical elements for constructing biomass. Pace-of-life theory posits that organisms trade-off allocation of these resources to assimilation, growth, and reproduction, with species falling on a slow-fast life history continuum (*1-4*). Modular organisms such as plants and social insects, however, organize the allocation of these tasks in a unique way. In modular organisms, resource acquisition and assimilation are specifically tasked to resource-acquiring modules that are non-reproductive (*5*). The evolution and optimisation of these modules can be described using economic concepts of investment and return of resources and nutrients (*6*). Such economic concepts have been widely developed in plants. For example, the Leaf Economics Spectrum (LES) scheme (*7*) underpins current understanding of plant investment strategies: plant species vary from investing in cheaply constructed ‘fast’ leaves with high nutrient concentrations of nitrogen and phosphorus, high metabolic rates, rapid assimilation of resources and short lifespans, to species with expensive but ‘slow’ leaves that are long-lived, with slow assimilation and metabolic rates.

It has been proposed that the leaf economic spectrum may also be applicable to other modular organisms such as social insects (*5*), but this hypothesis awaits testing. Ants represent a particularly complex form of social organisation where genetically related members of the colony act collectively as “superorganisms” to complete the functional tasks of a unitary organism (*8*). Under this social organisation, billions of individual ant workers gather and recycle the planet’s resources every day, performing essential ecosystem processes, from seed dispersal to nutrient cycling and predation (*9-12*). As for modular plants, this has been a highly successful ecological strategy with both ants and plants playing dominant roles in most terrestrial ecosystems (*13, 14*). Gibb et al. (*5*) hypothesised that ants and plants have converged on the same suite of approaches to acquire and distribute resources and that this could be indexed by a set of measurable traits. This leading hypothesis could unify the study of ant colonies with that of plants under the same economic spectrum umbrella, but it has not been empirically tested.

Here, we advance generality in economic theories of life by testing whether ant species fall along a fast-slow economic spectrum akin to the leaf economics spectrum. Sampling across extensive climatic and soil phosphorus gradients (fig. S1, table S1), we acquired trait data from individual workers collected from 123 species from seven subfamilies, including the five most diverse globally. Previous work to describe the resource economics of ant colonies has been fragmented and has failed to formally quantify the process of colony investment of nutrients and biomass in ant workers with a methodology that is scalable to large datasets (*15-19*). Here, we use eight traits to quantify the economic spectrum (*5*). Six traits directly parallel those used in the leaf economics spectrum: (1) Mass-specific standard metabolic rate, representing the *energy maintenance costs* of workers, as it does for plant leaves (*20*); (2) Mass density measured as mass:volume ratio and indicative of anatomical structure (e.g., cuticle thickness (*21*)), representing an *investment cost* equivalent to leaf-mass per area; (3) Nitrogen and (4) phosphorus concentrations of body tissue representing chemical *investment costs* (*22*); (5) Mass-specific assimilation rate (intake of proteins and carbohydrates per unit time per unit mass measured in laboratory feeding trials), representing *potential revenue* for the colony and equivalent to photosynthetic rate in plant leaves (*5*); (6) Relative worker lifespan (measured as days of survival in laboratory trials), equivalent to leaf lifespan, representing the *duration of the revenue stream* (*7*) that should be *optimised* to balance lifetime net resource costs and benefits (*23, 24*). Finally, we also include two ant-specific traits, (7) worker body mass, and a relative measure of (8) worker number (here referred to as colony size), that can be altered to change overall colony mass thereby influencing the total colony budget (*25*).

Chemical elements of nitrogen and phosphorus play slightly different roles in plant leaves and metazoan bodies, and we would expect a divergence in the co-variation of these two traits for plant and ant economic spectra. In leaves, concentrations of nitrogen and phosphorus are positively correlated, with high values of both indicating a fast economic strategy (*7*). In small-bodied animals like ants, however, nitrogen and phosphorus should trade-off (*26*). Nitrogen creates biological structure, such as costly cuticle, spines and dense muscles, while phosphorus is a critical component of the protein synthesis machinery of ribosomes (r-RNA) that sets the pace of worker growth (*27*). Broadly, greater allocation to nitrogen-rich structure is hypothesised to slow growth, while greater allocation to phosphorus rich r-RNA results in fast growth; hence these two elements should be negatively correlated in ants creating a fast-slow stoichiometric spectrum (*26*).

We present our predictions of how these eight traits interrelate to form a fast-slow economic spectrum and propose four key hypotheses that underlie these predictions (Fig. 1). These four hypotheses integrate across a portfolio of underlying ecological theory to support an economic spectrum in ants from first principles (*28*). Firstly, metabolic theories relate mass and energy use to allometric scaling laws that apply to all organisms (*3, 20*). Secondly, theories of ageing support the hypothesis that initial investment from the colony, resource acquisition rates, and extrinsic foraging risks are correlated to shape intrinsic lifespans of ant workers (*23, 29, 30*). Thirdly, pace of life theory posits a trade-off between offspring quality and quantity (*21, 31*), while social evolutionary theory suggests more complex colony organization reduces offspring reproductive potential (*32*): both theories link worker number (quantity) to worker structural biomass (size and quality). Fourthly, as outlined above, ecological stoichiometry theory suggests a trade-off in nitrogen and phosphorus allocation that should shape the pace of economic growth of the colony (*26*). Finally, we predict that the relative positions of ant species along the economic spectrum will be both phylogenetically conserved and modulated by the environment (Fig.1) and explore this by testing for evolutionary and biogeographic signatures.

**Fig. 1:**
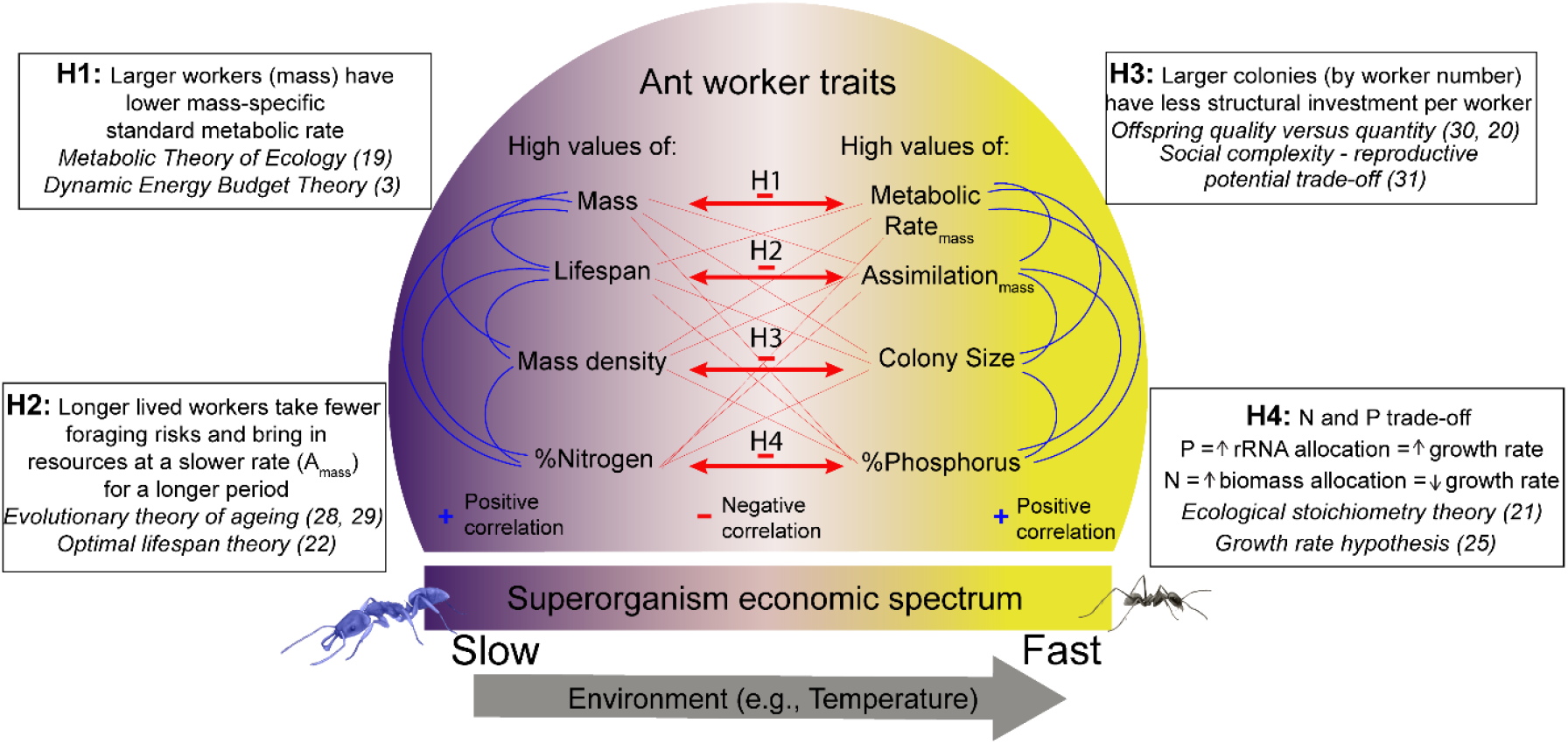
Conceptual framework of the hypothesised relationships between eight ant worker traits that together place species along a fast-slow economic spectrum. Predicted positive trait correlations shown by blue, predicted negative trait correlations shown in red. Four key trait trade-offs and their hypothesised relationships are highlighted and explained in text boxes H1, H2, H3, H4 with reference to underlying bodies of theory (see reference list: (*3, 20-23, 26, 29-32*)). High trait values of body mass, lifespan, mass density, and nitrogen concentration and low trait values for the other four traits are predicted to indicate a slow strategy. High trait values of mass-specific metabolic rate, mass-specific assimilation (resource intake) rate, colony size (worker number), and phosphorus concentration and low trait values for the other four traits are predicted to indicate a fast strategy. Species positions along the superorganism economic spectrum indicate the pace of investment and return of resources into ant workers and back to the colony. Species positions may be modulated by the abiotic and biotic environment, for example, ambient temperature could shift communities towards fast ecological strategies.

### The ant economic spectrum

We quantified relationships among the eight economic traits and decomposed the portion of trait correlation associated with phylogeny (conserved trait correlation *sensu* (*33*)) and that independent of phylogeny (non-phylogenetic) using multi response phylogenetic mixed models (*33, 34*). Our analyses revealed sets of phylogenetically conserved, co-varying traits that represent a clear trade-off in investment and resource return, forming an overall spectrum of fast-slow economic strategy (Fig. 2a, b). Species that invest more biomass and nitrogen in workers have small colonies of larger, longer-lived workers with low %P, lower maintenance costs (low mass-specific metabolic rate; H1, Fig. 1) and slow rates of resource return (low mass-specific assimilation rate) – an overall slow economic strategy (Fig. 2a, b). Species that invest less biomass and nitrogen into their workers have higher metabolic costs, high %P (representing fast growth rate; H4, Fig. 1) and quick rates of return on revenue invested (*20*). High metabolic rates likely increase the foraging tempo of ant workers, increasing the rate of resource uptake and thereby offsetting these high energy maintenance costs (*35*). These species with a fast economic strategy achieve larger colony sizes made up of smaller workers with less structural investment, shorter lives, and faster worker turnover (as predicted by H2, H3: Fig. 1) (Fig. 2a, b).

**Fig. 2:**
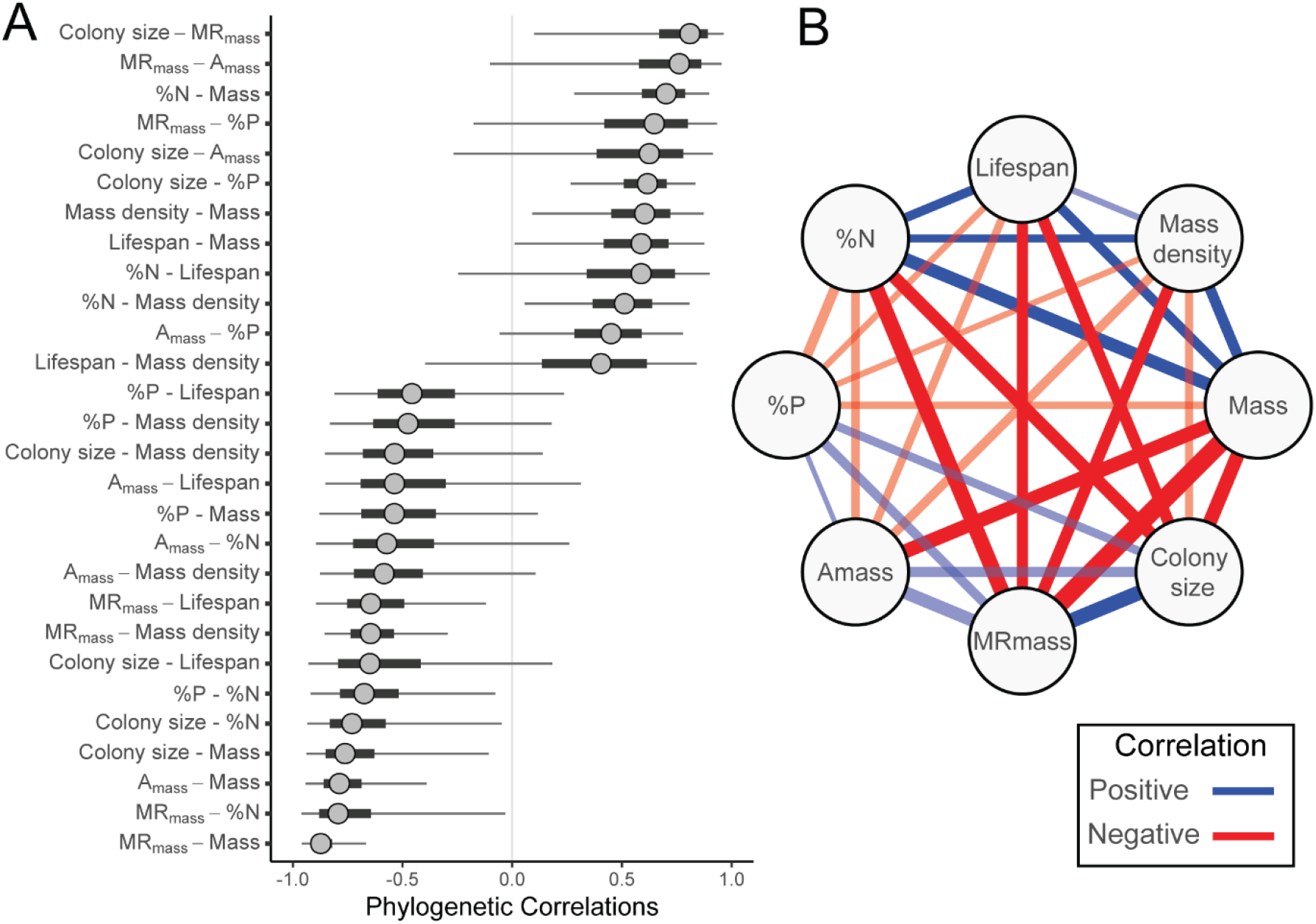
Two sets of correlated traits trade-off to create a fast-slow economic spectrum. (A) Posterior summaries of phylogenetic correlation coefficients from MR-PMM analysis, showing the posterior mean (point), 50% (heavy wick) and 95% (light wick) credible intervals. (B) Network diagram showing phylogenetic trait correlations significant at 95% (bold) and 70-90% (faded) credible interval. Blue lines = positive relationships, red lines = negative relationships. Line width indicates strength of correlation. Phylogenetic correlations are estimated with greater uncertainty than standard between-species correlations (*34*), meaning 95% CI are likely to be conservative. Despite some long-tailed posterior distributions, the absolute posterior mean estimates for each correlation coefficients were all >0.4, identifying an integrated trait network that strongly supports a fast-slow economic spectrum. A_mass_ = mass-specific resource assimilation, MR_mass_ = mass-specific metabolic rate.

Trait-trait phylogenetic correlations were strong (Fig. 2a, table S2) (majority >90% credible interval; Data S1). Two traits with less data coverage, %P and mass-specific assimilation rate showed weaker phylogenetic trait-trait correlations than other traits (Fig. 2a, table S2). Phylogeny-independent trait-trait correlations were much weaker overall for all trait pairs (table S2, fig. S2, S3, Data S1). Ant economic traits also had strong phylogenetic signal indicating conserved trait correlation (average posterior mean *λ* across traits: 0.62, range: 0.36 – 0.84; fig. S2, table S3). Lifespan (mean *λ* = 0.36) and nitrogen (mean *λ* = 0.41), however, were less phylogenetically conserved than other traits. Model validation confirmed that the fitted model generated plausible data for all response variables including for traits that had lower data coverage (posterior predictive checks: fig. S4). Leave-one-out cross validation indicated that model predictions were robust for predicting new data and were not overly influenced by any one species (LOO-CV: fig. S5).

Selection has therefore favoured workers that gather resources at a rate that reflects their investment costs and the duration of their revenue stream (*28, 29*). This parallels optimal lifespan theory for leaves (*23*), where average net resource carbon gain is maximized over the leaf life cycle (*36*). Consequently, some trait combinations will be rare or never observed (*37*). For example, tough leaves, or robust ant workers with short lifespans, will not have enough photosynthetic/foraging time to pay off high resource investment.

### Biophysical laws shape the economic spectrum

The independent component of trait-trait relationships (not associated with phylogenetic history) revealed three notable correlations that are likely to be driven by fundamental biophysical principles dictating energy-mass balance (*3, 20*). Mass-specific metabolic and assimilation rates showed an allometric negative relationship with body mass for both phylogenetic (Fig. 2a, b) and independent trait correlations (table S2, fig. S3). From cells to organisms to superorganisms (*38*), metabolic rate (*B*) scales allometrically with mass, according to a power function *B* = *a*M^β^ where, β is typically less than 1 (>-1 and <0 when mass-specific). Here, the mean slope of the independent component of these relationships was -0.55 (mass-metabolic) and -0.58 (mass-assimilation) respectively, indicating that smaller workers use more energy and consume food resources at a faster rate per unit body mass than do larger workers (as predicted in H1: Fig. 1, table S2).

Several theories have proposed that physio-chemical factors place fundamental mass-based limits on the metabolic and assimilation rates of organisms (*39*). These allometric relationships are hypothesised to relate to either nutrient supply networks (the basis of metabolic theory of ecology (*20*)) or to surface-area scaling dynamics of energy reserve pools and volume of structural biomass (dynamic energy budget theory (*3*)). The non-phylogenetic proportion of these correlations is therefore likely due to biophysical laws that determine the boundaries of economic-strategy space.

### Clades partition economic-strategy space

When visualized in multidimensional trait space, the ant economic spectrum was observed to occur over two axes of variation. The first axis represents the ant worker economic spectrum (AWES), explaining 61% of variance. This axis was characterised by the correlation between biomass investment and lifespan trading off against mass-specific metabolic rate, mass-specific assimilation rate, and colony size (Fig. 3). The second axis of variation represents a stoichiometry spectrum (SS), represented by the trade-off of N and P, explaining 13% of variance (as predicted by H4: Fig. 1). This shows a distinct departure from the leaf economics spectrum in which N and P in plant leaves are positively correlated and co-vary with other LES traits over one axis of variation (*7*). The N-P trade-off in ants aligns with ecological stoichiometry theory applicable to small metazoans (*27*). Although we did not explicitly test worker growth rates here, the negative association between N and P is consistent with our hypothesis that that biomass investment in nitrogen-rich structures and chemicals (e.g., thick cuticle (*21*), proteinaceous venoms (*40*)) is likely to be traded-off with worker growth rate represented by %P (*21, 31*) (Fig. 2b, Fig. 3).

**Fig. 3:**
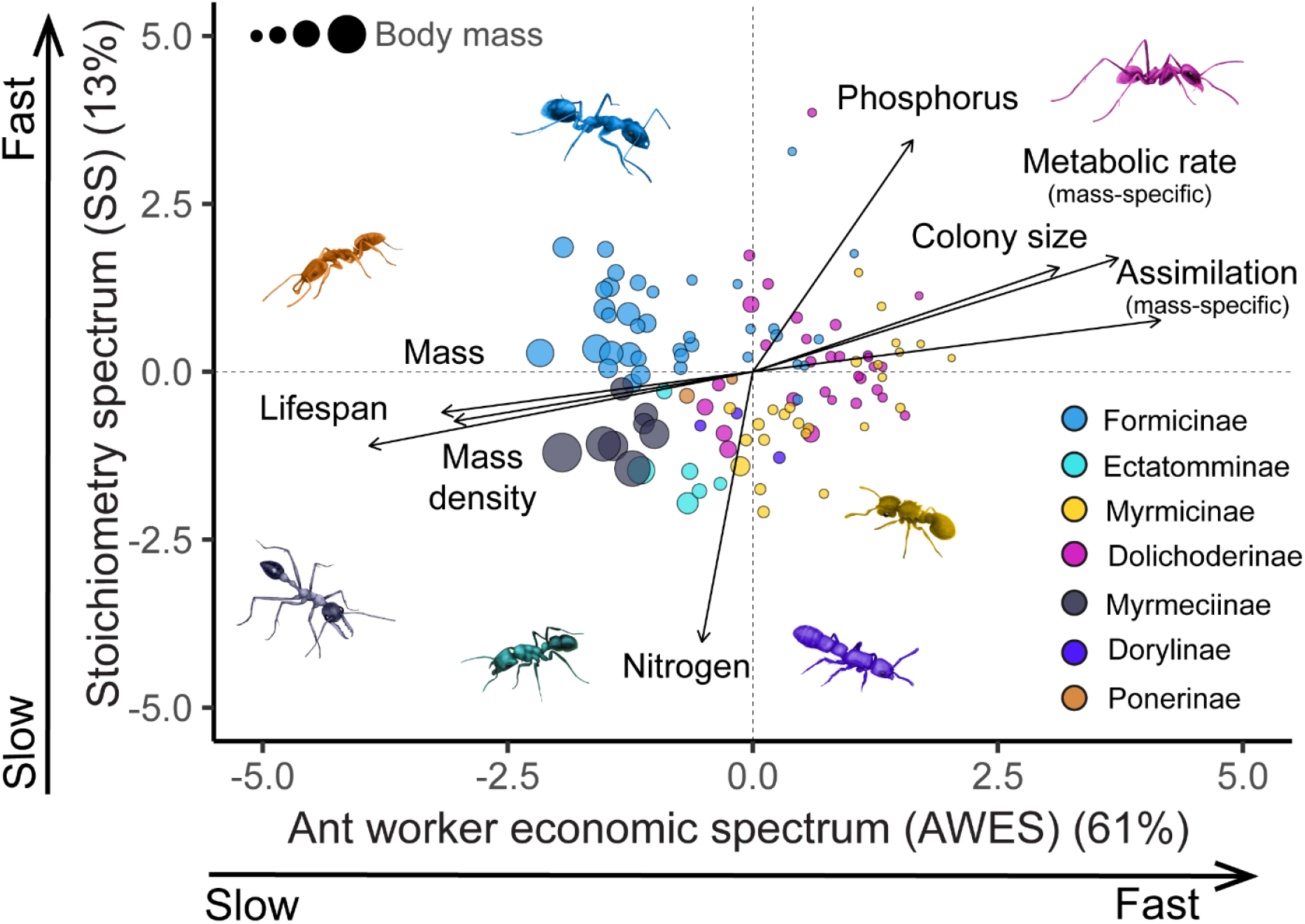
Mapping the superorganism economic spectrum onto multidimensional space. Showing varimax-rotated PCA from gap-filled species-averaged trait data from 305 colonies, with the 123 species as points and coloured as subfamily groups. Percentage variation explained in brackets for axis 1 (ant worker economic spectrum - AWES) and axis 2 (stoichiometry spectrum - SS). Size of points represents dry body mass scaled between 1-10. Ant icons are representative genera from each subfamily, anticlockwise from top left: Formicinae = *Camponotus*, Ponerinae = *Anochetus*, Myrmeciinae = *Myrmecia*, Ectatomminae = *Rhytidoponera*, Dorylinae = *Lioponera*, Myrmicinae = *Meranoplus*, Dolichoderinae = *Iridomyrmex*.

The different subfamilies occupy distinct regions of multidimensional economic strategy space (MANOVA – Axis 1: F_(6, 116)_ = 26.86, p < 0.0001, Axis 2: F_(6, 116)_ = 16.02, p < 0.0001, Fig. 3, 4). For example, two of the most globally diverse subfamilies, Dolichoderinae and Myrmicinae, generally occupy the fast end of the ant worker economic spectrum. Within that, myrmicines are placed on the slower end of the stoichiometry spectrum with higher N content compared with dolichoderines (Fig. 3, fig. S6). Many dolichoderine genera had a relatively fast strategy over all traits including the behaviourally dominant *Iridomyrmex* and *Anonychomyrma* (Fig. 4). Larger-bodied species of Formicinae, another globally diverse clade, fall on the slower end of the ant worker economic spectrum, but on the fast end of stoichiometry spectrum, with high phosphorus and low nitrogen concentrations, likely due to lower investment in cuticle thickness and a lack of proteinaceous venoms (*21, 40*) (Fig. 3, Fig. 4). Meanwhile, the more densely built Ectatomminae and Myrmeciinae (*41*) exhibit a slow strategy across both axes (Fig. 3, 4, fig. S6). This partitioning of economic spectrum space across the two axes indicates that there are different templates for being a “slow” or “fast” ant (Fig. 4).

**Fig. 4:**
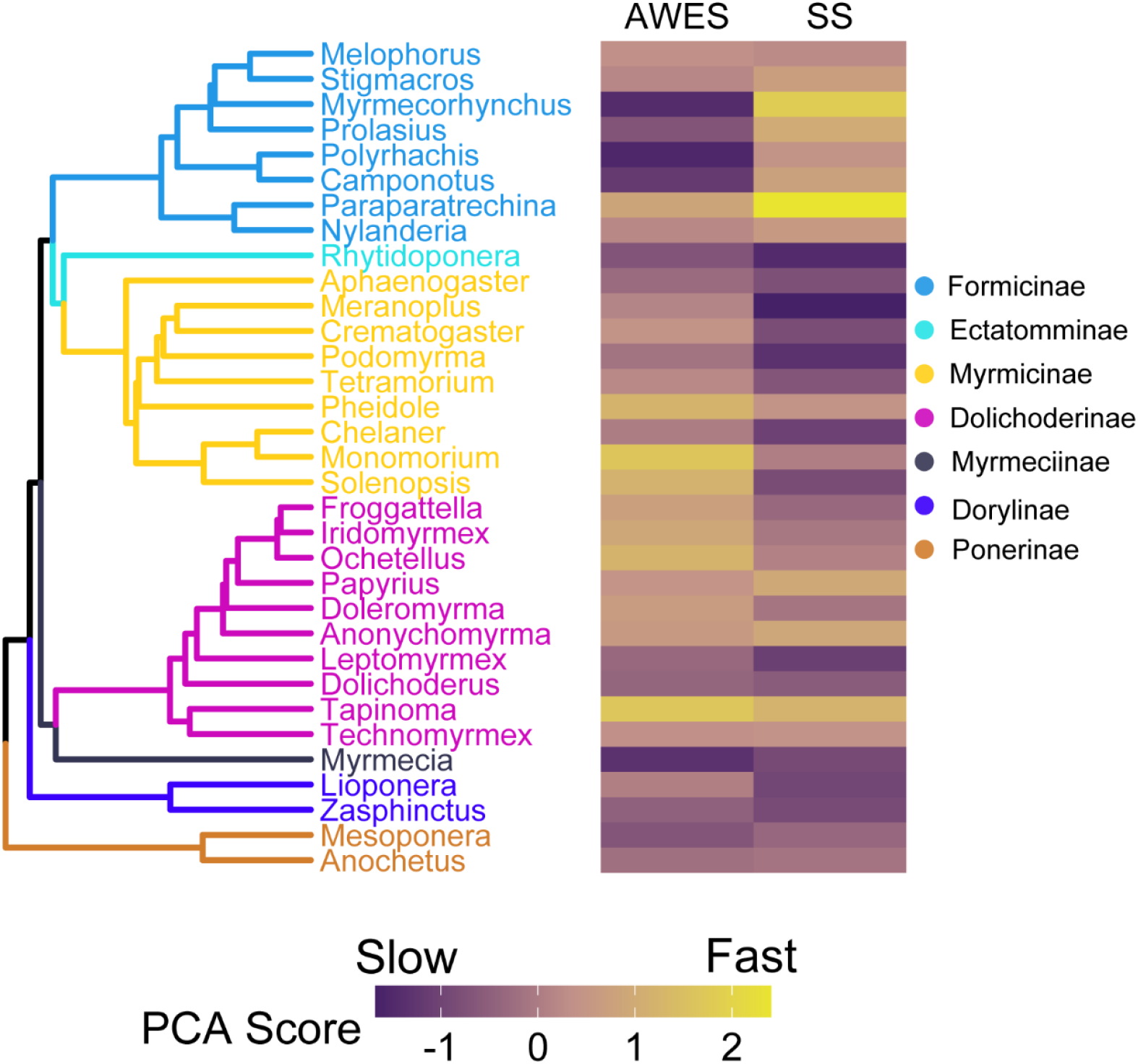
Mapping the superorganism economic spectrum onto the ant phylogenetic tree at the genus level. Showing PCA scores from Fig. 3. For axis 1: Ant Worker Economic Spectrum (AWES) and for axis 2: Stoichiometry Spectrum (SS). Slow values (darker colours) for AWES signify large body mass, long lifespan, high mass density, low mass-specific metabolic and assimilation rates and small colony sizes. Fast values (lighter colours) represent the opposite values for each trait. Slow values (darker colours) for stoichiometry spectrum (SS) represent high nitrogen concentration and low phosphorus concentration, with the opposite ratio for fast ecological strategies (lighter colours). This demonstrates that niche partitioning is occurring along two axes of the ant economic spectrum (see PCA Fig. 3 to see these patterns mapped onto multidimensional space, see fig. S6 for subfamily tree). Note, genus *Arnoldius* scored very high on the SS spectrum and is removed in this figure to improve visualization of colour gradient, see fig. S7 for figure that includes *Arnoldius*.

### Full economic spectrum persists in all environments

Other studies have shown that some ant traits obey universal physiological constraints (*38, 42*) and are strongly phylogenetically conserved (*43*), while others vary across both geographic (*25*) and regional environmental gradients (*44, 45*). Here, however, we found that the full suite of economic strategies was present in all environmental conditions tested (fig. S8, table S4). Local-scale climate or soil phosphorus, for example, did not shape key traits linked to economic strategies (MR-PMM trait:fixed effects, average p-value = 0.7, Data S2). This is consistent with other studies showing that local communities can contain the entire regional pool of ant trait variation (*46*). Similar global patterns are found in plants: some leaf traits have relatively strong relationships with climate (e.g., leaf mass per area with aridity (*47*)) but species positions on the slow-fast economic spectrum appear largely independent of climate (*7*).

We further note that economic strategies can vary within-species at local scales through phenotypic plasticity. For instance, common garden experiments have revealed that exposure to predators can promote colony defensive allocation (e.g., to major castes (*48*)) and thus likely caste-specific stoichiometric allocation demands (*49*). Further, resource manipulation studies have shown that overall ant activity can increase with phosphorus supplementation (*50*). Thus, core features of the economic spectrum are presumably subject to environmental modulation at the population level. Here, it appears that species with both slow and fast strategies persist under a broad range of environmental conditions. Future studies could investigate whether the relative abundances of species differing in economic spectrum positions changes with the environment (e.g. through community weighted means), and particularly with respect to disturbance regimes, which have previously been linked with the key trait of body size at a global scale (*51, 52*).

## Conclusion

We have described a phylogenetically conserved economic spectrum for a globally dominant group of superorganisms. Based on species presence, the spectrum did not vary systematically with climate and environmental gradients. Our results suggest that early lineages of ants likely diverged in their economic strategies and subsequently diversified (*53*), thereby locking in these fundamental templates of investment and resource return. We have presented a foundational test of this hypothesis using Australian ant species. We call for more global sampling, for example, comparing economic spectrum traits in ants across broader biogeographic gradients, such as temperate versus tropical, and also in other social or modular organisms. Given the strong role of evolutionary history, compared with climate and environment, in the economic spectrum, we predict that more globally extensive sampling of ant communities would align with our key results.

The economic spectrum framework has clear potential for modelling the resource economics of other social insects who likewise invest in their non-reproductive workers to bring in resources that promote colony fitness. Workers of social insect colonies gather resources across vast scales and have major impacts on habitats across our planet. Globally, ants can increase crop yields through their positive role on soil health (*12*), managed bee colonies pollinate one third of our crops (*54*) and termites decompose more than half of the deadwood in tropical forests (*9*). In plants, aggregated economic spectrum trait values of communities, together with biotic species interactions, correlate with ecosystem productivity, directly linking the scaling of resource economics from leaves to ecosystems (*55*). Mapping social insects along the fast-slow worker economic spectrum could be similarly used to understand the link between rates of resource use and the allocation of energy and nutrients from colony to community scales. Economic trait values of local assemblages could further be combined with data on ecological roles to estimate the rates of ecosystem services such as decomposition or pollination (*51*).

A major goal of ecology is to identify broadly applicable principles that provide a mechanistic understanding of how living things behave and interact with their environment. Our findings show that evolutionary optimisation and the biophysical laws of energy and mass balance have jointly shaped the economics of superorganisms such as ants in similar ways to plants indicating common solutions to resource allocation problems across the tree of life.

## Acknowledgements

We thank and acknowledge the Wurundjeri people of the Kulin Nation, the Wotjobaluk, the Ngiyampaa, the Durramurragal, the Gubbi Gubbi, the Boonwurung, Bunurong, and Gunaikurnai people on whose lands this field work was conducted. We thank Emily House for access to Glen Echo. Field work was carried out under Permit SL102675 NSW Department of Planning, Industry and Environment, Permit AA-0000328 Parks Victoria, and under animal ethics committee permit AEC22001. We thank Luke A. Yates for advice and support with statistical modelling. We thank Ian J. Hammer for technical assistance with metabolic rate assays. We thank Wednesday Elgar, Stephanie Nicholls, Andrew Chalmers, Hannah Elmes, Nicholas Gale, Lia Herrera Grau and volunteers for helping with field and lab work.

## Funding

Australian Research Council Discovery Project DP210101630 (HG, IJW, NJS, CLP, TRB)

Australian Government RTPS PhD Funding (HLR)

Australian Research Council Discovery Project DP190100341 (SLC)

Human Frontiers of Science Program grant RGEC26/2024 (TRB)

The Royal Society RGS\R1\221207 (TRB)

## References

1. K. J. Mathot, W. E. Frankenhuis, Models of pace-of-life syndromes (POLS): a systematic review. Behavioral Ecology and Sociobiology 72, 41 (2018).

2. J. R. Burger, C. Hou, C. A. Hall, J. H. Brown, Universal rules of life: metabolic rates, biological times and the equal fitness paradigm. Ecology Letters 24, 1262–1281 (2021).

3. S. A. L. M. Kooijman, Dynamic energy budget theory for metabolic organisation. (Cambridge university press, 2010).

4. C. Zhang, I. J. Wright, U. N. Nielsen, S. Geisen, M. Liu, Linking nematodes and ecosystem function: a trait-based framework. Trends in Ecology & Evolution, (2024).

5. H. Gibb et al., Ecological strategies of (pl)ants: Towards a world-wide worker economic spectrum for ants. Functional Ecology 37, 13–25 (2023).

6. R. R. Junker et al., Towards an animal economics spectrum for ecosystem research. Functional Ecology 37, 57–72 (2022).

7. I. J. Wright et al., The worldwide leaf economics spectrum. Nature 428, 821–827 (2004).

8. J. J. Boomsma, R. Gawne, Superorganismality and caste differentiation as points of no return: how the major evolutionary transitions were lost in translation. Biological Reviews 93, 28–54 (2018).

9. H. M. Griffiths, L. A. Ashton, T. A. Evans, C. L. Parr, P. Eggleton, Termites can decompose more than half of deadwood in tropical rainforest. Current Biology 29, 118–119 (2019).

10. H. M. Griffiths et al., Ants are the major agents of resource removal from tropical rainforests. Journal of Animal Ecology 87, 293–300 (2018).

11. A.-M. Klein et al., Importance of pollinators in changing landscapes for world crops. Proceedings of the Royal Society B: Biological Sciences 274, 303–313 (2007).

12. D. Wu, E. Du, N. Eisenhauer, J. Mathieu, C. Chu, Global engineering effects of soil invertebrates on ecosystem functions. Nature, (2025).

13. J. M. Kass et al., The global distribution of known and undiscovered ant biodiversity. Science Advances 8, eabp9908 (2022).

14. S. Díaz et al., The global spectrum of plant form and function. Nature 529, 167–171 (2016).

15. W. S. Creighton, The ants of North America. Bulletin of the Museum of Comparative Zoology 104, 1–585 (1950).

16. W. M. Wheeler, Ants: their structure, development and behavior. (Columbia University Press, New York, 1910), pp. 663.

17. G. Oster, E. Wilson, Caste and ecology in the social insects. (Princteon University Press, Princeton, NJ, 1978), pp. 352.

18. P. Calabi, S. D. Porter, Worker longevity in the fire ant Solenopsis invicta: ergonomic considerations of correlations between temperature, size and metabolic rates. Journal of Insect Physiology 35, 643–649 (1989).

19. M. Kaspari, M. M. Byrne, Caste allocation in litter Pheidole: lessons from plant defense theory. Behavioral Ecology and Sociobiology 37, 255–263 (1995).

20. J. H. Brown, J. F. Gillooly, A. P. Allen, V. M. Savage, G. B. West, Toward a metabolic theory of ecology. Ecology 85, 1771–1789 (2004).

21. C. Peeters, M. Molet, C.-C. Lin, J. Billen, Evolution of cheaper workers in ants: a comparative study of exoskeleton thickness. Biological Journal of the Linnean Society 121, 556–563 (2017).

22. J. J. Elser et al., Biological stoichiometry from genes to ecosystems. Ecology Letters 3, 540–550 (2000).

23. K. Kikuzawa, A cost-benefit analysis of leaf habit and leaf longevity of trees and their geographical pattern. The American Naturalist 138, 1250–1263 (1991).

24. B. H. Kramer, G. S. van Doorn, B. M. S. Arani, I. D. Pen, Eusociality and the Evolution of Aging in Superorganisms. American Naturalist 200, 63–80 (2022).

25. M. Kaspari, Global energy gradients and size in colonial organisms: worker mass and worker number in ant colonies. Proceedings of the National Academy of Sciences 102, 5079–5083 (2005).

26. J. J. Elser et al., Growth rate–stoichiometry couplings in diverse biota. Ecology Letters 6, 936–943 (2003).

27. J. F. Gillooly et al., The metabolic basis of whole-organism RNA and phosphorus content. Proceedings of the National Academy of Sciences 102, 11923–11927 (2005).

28. P. A. Marquet et al., Scaling and power-laws in ecological systems. Journal of Experimental Biology 208, 1749–1769 (2005).

29. L. Keller, M. Genoud, Extraordinary lifespans in ants: a test of evolutionary theories of ageing. Nature 389, 958–960 (1997).

30. N. J. Lemanski, N. H. Fefferman, How life history shapes optimal patterns of senescence: implications from individuals to societies. The American Naturalist 191, 756–766 (2018).

31. E. R. Pianka, On r-and K-selection. The american naturalist 104, 592–597 (1970).

32. Bourke, Colony size, social complexity and reproductive conflict in social insects. Journal of Evolutionary Biology 12, 245–257 (1999).

33. M. Westoby, L. Yates, B. Holland, B. Halliwell, Phylogenetically conservative trait correlation: Quantification and interpretation. Journal of Ecology 111, 2105–2117 (2023).

34. B. Halliwell, B. R. Holland, L. A. Yates, Multi-response phylogenetic mixed models: concepts and application. Biological Reviews 100, 1294–1316 (2025).

35. K. S. Mason, C. L. Kwapich, W. R. Tschinkel, Respiration, worker body size, tempo and activity in whole colonies of ants. Physiological Entomology 40, 149–165 (2015).

36. H. Wang et al., Leaf economics fundamentals explained by optimality principles. Science Advances 9, eadd5667 (2023).

37. M. Castorena, M. E. Olson, B. J. Enquist, A. Fajardo, Toward a general theory of plant carbon economics. Trends in Ecology & Evolution 37, 829–837 (2022).

38. J. Z. Shik, C. Hou, A. Kay, M. Kaspari, J. F. Gillooly, Towards a general life-history model of the superorganism: predicting the survival, growth and reproduction of ant societies. Biology Letters 8, 1059–1062 (2012).

39. J. L. Maino, M. R. Kearney, R. M. Nisbet, S. A. Kooijman, Reconciling theories for metabolic scaling. Journal of Animal Ecology 83, 20–29 (2014).

40. B. D. Blanchard, C. S. Moreau, Defensive traits exhibit an evolutionary trade-off and drive diversification in ants. Evolution 71, 315–328 (2017).

41. J. T. Buxton, K. A. Robert, A. T. Marshall, T. L. Dutka, H. Gibb, A cross-species test of the function of cuticular traits in ants (Hymenoptera: Formicidae). Myrmecological News 31, 31–46 (2021).

42. C. Hou, M. Kaspari, H. B. Vander Zanden, J. F. Gillooly, Energetic basis of colonial living in social insects. Proceedings of the National Academy of Sciences 107, 3634–3638 (2010).

43. J. Vizueta et al., Adaptive radiation and social evolution of the ants. Cell, (2025).

44. J. Z. Shik, X. Arnan, C. S. Oms, X. Cerdá, R. Boulay, Evidence for locally adaptive metabolic rates among ant populations along an elevational gradient. Journal of Animal Ecology 88, 1240–1249 (2019).

45. L. Leahy, B. R. Scheffers, S. E. Williams, A. N. Andersen, Arboreality drives heat tolerance while elevation drives cold tolerance in tropical rainforest ants. Ecology 103, e03549 (2022).

46. M. Kaspari, N. A. Clay, J. Lucas, S. P. Yanoviak, A. Kay, Thermal adaptation generates a diversity of thermal limits in a rainforest ant community. Global Change Biology 21, 1092–1102 (2015).

47. Ü. Niinemets, Global-scale climatic controls of leaf dry mass per area, density, and thickness in trees and shrubs. Ecology 82, 453–469 (2001).

48. A. S. Yang, C. H. Martin, H. F. Nijhout, Geographic variation of caste structure among ant populations. Current Biology 14, 514–519 (2004).

49. A. D. Kay, S. Rostampour, R. W. Sterner, Ant Stoichiometry: Elemental Homeostasis in Stage-Structured Colonies. Functional Ecology 20, 1037–1044 (2006).

50. J. Bujan, S. J. Wright, M. Kaspari, Biogeochemical drivers of Neotropical ant activity and diversity. Ecosphere 7, e01597 (2016).

51. F. de Bello et al., Raunkiæran shortfalls: Challenges and perspectives in trait-based ecology. Ecological monographs 95, e70018 (2025).

52. H. Gibb et al., Habitat disturbance selects against both small and large species across varying climates. Ecography 41, 1184–1193 (2018).

53. C. S. Moreau, C. D. Bell, R. Vila, S. B. Archibald, N. E. Pierce, Phylogeny of the Ants: Diversification in the Age of Angiosperms. Science 312, 101–104 (2006).

54. FAOSTAT, ProdSTAT Database. 2022 (https://www.fao.org/faostat/en/#data/QCL Accessed 28-09-2022).

55. P. B. Reich, The world-wide ‘fast–slow’plant economics spectrum: a traits manifesto. Journal of ecology 102, 275–301 (2014).

