## Supplementary material for "The economic strategies of superorganisms": Methods and Supplementary Material

### Supplementary Material
### The economic strategies of superorganisms

**Authors:** Lily Leahy[1*], Hannah L. Riskas[1], Ben Halliwell[2,3], Ian J. Wright[4,5], Amelia G. Carlesso[1,6], Nathan J. Sanders[7], Tom R. Bishop[8,9], Catherine L. Parr[9,10], Steven L. Chown[11], Jonathan Z. Shik[12], Alan N. Andersen[13], Heloise Gibb[6, 14]

#### Materials and Methods

Field locations

Ants were sampled from six locations along the east coast and inland of south-eastern Australia (fig. S1) representing a gradient of low to high aridity (421 – 1283 mm annual precipitation) and temperature (13 – 20 ˚C mean annual temperature). Locations were each sampled over a one-week period between June 2022 to April 2023. Within each location, two sites with contrasting soil phosphorus status were chosen using the CSIRO Soil and Landscape Grid National Soil Attribute Map – Total Phosphorus (3" resolution) release 1.V5 (*1*). Within each high P and low P site, four plots (10 x 10 m) separated by ~200 m, were established (total of eight plots per site). Soil phosphorus status of the plots was confirmed using soil chemical analysis. Nine soil cores (~5 cm diameter, 15 cm deep) were taken per plot, combined, and air-dried giving one soil sample per plot. Soil P was assessed through soil chemical analysis of Total P (17C1 Aqua regia block digestion; (*2*)) at Environmental Analysis Laboratory, Southern Cross University.

Ant field sampling

Each plot was comprehensively sampled for ants for two, one-hour blocks over two days such that each plot was surveyed in both the morning (~0900 – 1300 hours) and afternoon (1300 – 1700 hours). A final day of sampling was undertaken targeting specific plots to increase the number of species and colony replicates. Ten baits in 5 ml plastic vials, five with tuna, five with honey, were placed on the ground along a 50 m transect at the beginning of the search to attract foraging ants. Ants were collected from baits if they had formed a clear foraging trail which indicated the individuals were from the same colony. Hand searches involved locating nests by checking under rocks, logs, tussocks, coarse woody debris, and vegetation and looking for ant foraging trails. Ants foraging arboreally in columns were assumed to be from the same colony and were sampled using aspirators or brushing ants into a container with a paint brush. To lure ants from nests that were difficult to access, another five to ten traps of each bait type were placed at nest entrances.

Climate data

We modelled microclimate for the sites using the *micro_ncep* function in package NicheMap R for the years 2009-2023 (*3*). This function uses the *microclima* package (*4*) as well as the RNCEP (*5*) and elevatr (*6*) packages to connect to the 6-hourly 2.5 x 2.5-degree gridded historical NCEP data (global scope). It then downscales climate to hourly estimates accounting for local terrain effects (~30 x 30 m) including elevation-induced lapse rates, coastal influences and cold-air drainage (*3*). We parametrised the model for each site using shade values which were obtained from field vegetation surveys. For each plot, nine 50 $cm^2$ quadrats were placed at random and % cover of canopy, understorey, and shrub layer (where present) was recorded. The values were summed with the maximum value set at 100% cover. The values were then averaged per plot and then per site to provide a shade value. We used a single shade value per site-model. Default values were used for other parameters in the model. We extracted temperature (T) and relative humidity (RH) at 1.2 m standard height above ground and averaged these values to get daily averages and the annual average across all 15 years. We used T to calculate saturation water vapor pressure ($e_s$) and RH to calculate VPD (*7*) as:

$$e_s = 0.61078\exp\left(\frac{17.27T}{T + 237.3}\right)$$

$$VPD = e_s - \left(\frac{(RH)(e_s)}{100}\right)$$

Ant sampling

We aimed to collect as many ant species as possible from each site over three days of sampling. During sampling, live worker ants were collected either from foraging trails or nests using hand searches and baiting. We aimed to collect a minimum of 30 workers (and up to ~200 individuals) per colony, taken from 1-7 colony replicates per species per site (average of 2 colony replicates per species per site). In the field, ants were allocated to morphospecies within genera or species complexes based on morphological observations using a microscope. Following field collection, 3-5 individuals from each colony were pinned and assigned a species or a species complex. Voucher specimens were deposited in the CSIRO TERC collection in Darwin, Australia, and the Gibb Lab collection at La Trobe University, Australia. Live field collected colonies were placed into plastic containers with the top 10 cm of the inner rim lined with fluon, given a honey-water mix (50:50) (all ant workers consume some carbohydrates in their diet: (*8*)) and hydration via a saturated cotton ball every two days and housed at 20˚C. Ants were transported back to the lab for trait measurements within 1-4 days from collection. Brood, alates, and major workers were collected but were not used for any trait analyses reported here. For polymorphic species (e.g. *Camponotus* or *Pheidole* species), we exclusively measured traits for minor workers.

Economic traits

*Metabolic rate*

Ants were housed as above while waiting to be trialled. Time between field collection and metabolic trials across sites ranged between 3-14 days. As digestion can influence metabolic rate, ants were starved but provided with water for 48 hours prior to trials. Carbon dioxide production ($VCO_2$) was used as a proxy for metabolic rate and was measured at a consistent temperature of 22˚C of using 8 Sable Systems International (SSI) multiple animal versatile energetics (MAVEn (SSI, Las Vegas, Nevada, USA)) systems, each attached to a Li-Cor 7000 $CO_2$/$H_2O$ infrared gas analyser (Li-Cor, Lincoln, Nebraska, USA), housed in a Panasonic MIR 352H-PE climate control cabinet (Panasonic Healthcare, Sakata, Japan). Ants were placed in sets of 10 individual 2 ml or 3 ml curettes (depending on the size of the individual) with airstream humidity maintained at 90.34 ± 0.6% RH to avoid desiccation. Each ant was measured twice for a period of 10 minutes each time with a 5-minute baseline in between individuals to account for drift in the Li-Cor 7000 measurements. An additional 20 minutes was added at the beginning (for settling) and the end (as contingency). The activity of each ant was measured simultaneously as a unitless measurement using infrared light detectors in the MAVEn activity board.

Data treatment and extraction were conducted with the software Expedata (SSI). First $VCO_2$ data was corrected to standard temperature and pressure for a push system according to the equations of (*9*) and then baseline corrected using the Catmull-Rom spline method. To obtain standard metabolic rate (SMR) we then took the average of the lowest one minute of $VCO_2$ for each ant. $VCO_2$ was then inspected for outliers. Individuals with very high activity in the context of their respective colonies and species were removed (seven individuals). Individuals with erroneous values due to technical errors (e.g., temporary issues with flow rate) were removed (n = 103 individuals). We then systematically removed outliers that were 1.5 times the IQR of the data for each colony respectively (n = 184 individuals) as these could represent stressed or dying individuals. Finally, colonies with < 3 individuals remaining after these steps were removed from the dataset (n = 15 individuals). Together the outliers represented 9.56% of the original data resulting in a final dataset of 2805 individuals of 214 colonies of 114 species.

To explore whether activity during assays affected metabolic rate, we modelled $VCO_2$ as a function of the activity variables in a linear mixed effects models with species, colony ID, and survey location as random effects using the *lmer* function (*10*). There was no significant relationship between total activity over the assay and $VCO_2$, with random effects explaining all the variation in the model ($F = 0.14_{(1, 2718)}$, $p = 0.740$, $R_m^2 = 0.0002$, $R_c^2 = 0.72$). There was a significant relationship between activity at time of $VCO_2$ trace (slope of the absolute difference sum transformed activity (Slope ADS)) and $VCO_2$ (slope ± 95% CI: 0.54 (0.13 – 0.95), p = 0.01), but the fixed effect (Slope ADS) explained a very small amount of variance in the model ($R_m^2 = 0.0014$, $R_c^2 = 0.72$). Therefore, we did not include individual activity during assays as a covariate in downstream analyses. Minimum $VCO_2$ ($\mu L/hr^{-1}$) was then averaged per colony and converted to microwatts following equations from Chown, Gibbs, Hetz, Klok, Lighton and Marais (*11*): 1) Convert to Liters of $CO_2$, 2) Convert to kJ/hr (L CO2/hr x 24.65), 3) Convert from kJ/hr to Watts [kJ/hr x 0.2777], 4) Convert from Watts to microwatts [Watt x $10^6$].

*Lifespan*

Relative worker lifespan was measured in the lab using a subset of species collected from each of the six field locations. Following transport from the field to the lab, thirty workers per colony were transferred to new plastic containers as a colony fragment and housed in a controlled temperature room at 25˚C with humidity set to room ambient conditions. To simulate nest conditions and reduce stress, no light was provided except during feeding and hydration periods. Colony fragments (without brood or queens) were placed in a randomized position on the shelves to remove any location-specific effects of the controlled temperature room environment. They were provided with a diet of honey-water mix (50:50) and dried insects fed *ad libitum*, with hydration provided via a saturated cotton ball every two days. We monitored colony fragments daily to record and remove deceased ants that were placed in 5 ml tubes and instantly frozen at -20 °C. We ignored initial loss due to stress and began death counts 2 days after transfer to the controlled temperature room. Median survival time (in days) per colony was calculated using survival curves fitted with the Kaplan-Meier estimate using the function *survfit* in package "survival" (version 3.5-5) (*12*). We note two caveats: the age of individual workers collected from the field is unknown, further, lab lifespan of cohorts of workers without a queen can be biased and may not represent lifespan in the field realistically. We note that a previous study using the same experimental approach showed that field and laboratory measures of worker lifespan are positively correlated (*40*), lending support to the validity of our method. However, we suggest that laboratory measured lifespan represents a relative rather than absolute measure of worker lifespan.

*Body mass*

Field collected colonies which were not allocated to either metabolic rate assays or lifespan assays were immediately frozen at -20 °C upon collection from the field. All frozen ants (from metabolic assays, lifespan assays, and directly from the field) were dried at 50 °C for 48 hours. We then measured dry mass using a 0.001 mg precision XS3DU microbalance (Mettler Toledo). For ants from metabolic and lifespan assays, dry mass was measured for each individual and the average per colony calculated, for the remaining colonies, dry mass was measured for 10 ants per colony and their average mass calculated.

*Mass density*

We calculated mass density of worker ants as density = mass/volume. Head volume was used as a proxy for body volume. Head volume was calculated using the formula for the volume of an ellipsoid as $V = 4/3\pi abc$, where a = head width (HW), b = head length (HL), and c = head height (HH). The heads of dried ants were removed and placed under a microscope to measure HW and HL of five to ten individuals per colony. Measurements were taken in mm using a Leica microscope camera and Leica Application Suite (LAS version 1.4). We note that ant heads are unlikely to shrink in volume with drying due to their thick cuticles (i.e. in comparison to softer bodied insects). Head height was difficult to measure at scale using this technique. We predicted that HH would be a proportion of HW, but this proportion would vary by genus due to different morphology (note head shape is likely to be constrained at genus level and is therefore unlikely to show much variation amongst species within a genus (*13*)). We used front and side profile photos of pinned ants obtained from AntWeb (*14*) to calculate HH:HW as a proportion for each genus (n = 34) in the study. Using pinned AntWeb photos and imaging software Image J FIJI ver.1.54f (*15*) we measured HH and HW of 1-3 pinned individuals of three representative species (species were matched to the species in our dataset where possible) except for *Anochetus*, *Mesoponera*, and *Froggattella* which were represented by two species and *Paraparatrechina* which was represented by one species. We then calculated average HH:HW for each genus as a proportion between 0-1. HH:HW averaged 0.68 (± 0.08 SD) and ranged from 0.49 – 0.90. We then used the HH:HW proportion value per genus and the HW measurements from our dataset to estimate HH. Mass density of workers was then calculated as an average value per colony with units as $mg/mm^3$.

*Nitrogen and phosphorus body concentrations*

Dried ants (including removed heads) were combined per colony to undergo chemical analyses of nitrogen (%N) and phosphorus (%P). Gasters were removed prior to %N analysis as samples were simultaneously analysed for $\delta^{15}N$ which requires removal of recently consumed food located in the gaster (*16*) ($\delta^{15}N$ data not included in this study). Ant samples were analysed for nitrogen content (%N) using the EA (Dumas) method performed using an Isoprime (Micromass, Wythenshawe, Manchester, U.K.) with a Carlo Erba CE1100 elemental analyser (Fisons, Milan, Italy) at the Isotope Facility of the Farquhar Laboratory of the Research School of Biology, Australian National University, ACT, Australia. Approximately 1 to 2 mg (to 3 µg precision) of each dried sample was weighed into a tin capsule. The Isoprime corrects for drift and time variable source effects using $CO_2$ and $N_2$ reference pulses. Post processing corrections were made using laboratory standard materials (cane sugar, beet sugar, cysteine and glycine). Inhouse software (SecondRat) was used to assess the results and correct to the standards. Ant samples were analysed for phosphorus content (%P) using 17C1 Aqua regia block digestion (*2*) conducted at the Environmental Analysis Laboratory, Southern Cross University. Dried ants were ground and ~100.0 mg of each sample was weighed and analysed. Test and reference samples were used to correct for any drift or carry-over in the instrument. The references were calibrated for total %P.

*Colony size*

To estimate a relative measure of worker number (here referred to as colony size) we undertook a mark-recapture study on ant colonies in the field. At one field location (Nangak Tamboree Wildlife Sanctuary, Victoria) we sampled 17 species from 5 subfamilies and 10 genera, encompassing 34 colonies (2 colony replicates per species). Mark-recapture and field observations were conducted over several field sessions during the warmer months (February-April 2022, October-December 2022, January-April 2023). We selected cohorts of 30 to 100 foraging worker ants outside the nest per colony, prioritizing similarity in worker size to minimise the impact of polymorphic body sizes. We marked workers on their gasters using coloured paint marker pens. Every three months we marked a new cohort for each colony with a new colour. Within a cohort, individual ants were indistinguishable by their markings. We conducted biweekly hour-long observations, pausing only during heavy rain, until a month passed without sighting any marked workers. In each session, we documented two primary variables: the total number of foraging ants and the count of marked workers. Observations were made between 8:00 to 19:00 hours depending on when each colony was active. Activity times of the ants were based on previous field observations at this location (*17*).

To estimate colony size for each species, we employed Chapman's estimator (*18*), a refinement of the Lincoln-Petersen estimator, using mark-recapture data. We excluded only the days where no ants were observed. Chapman's estimator was calculated for each coloured cohort per colony and for each observation day using the formula:

$$\widehat{N}_C = \frac{(M-1)(C-1)}{R+1} - 1$$

where M is the total number of initially marked ants, C is the total number of ants observed on a recapture day, and R is the number of marked ants recaptured on a recapture day. We averaged these estimates of cohort size to derive a single colony size estimate.

*Assimilation rate*

We calculated assimilation rate for ant species from one field location (Nangak Tamboree Wildlife Sanctuary). This was measured in the lab as part of a larger nutritional geometry study on the trophic position of ants. We collected 30 individual workers from the same field colonies for which colony size was estimated above (n = 17 species, 34 colonies) over one week in March 2023. We transferred ants to plastic containers as a colony fragment and housed them in a controlled temperature room at 25 ˚C with humidity set to room ambient conditions. To simulate nest conditions and reduce stress no light was provided except during feeding and hydration periods. We formulated three dietary options, each moulded into a cube, giving different protein-to-carbohydrate (P:C) ratios: 1:3, 1:1, and 3:1. The diet cubes were prepared using a standardized protocol to ensure consistency (*19*). The experiment was conducted over six consecutive days, with each 24-hour interval marking a measurement cycle. At the end of each period, the diet cubes were weighed in their wet state, then dried for 48 hours at 60 ˚C and re-weighed to obtain a dry weight. We calculated the total intake values combining carbohydrate and protein intake over a six-day period. This rate was then divided by the number of workers in the colony fragment to give an assimilation rate as mg food consumed per worker per day.

*Trait data coverage and treatment*
Trait coverage was uneven across species (n = 123), genera (n = 34), and subfamilies (n = 7) (table S1). All traits were logged and scaled prior to analysis. Metabolic rate and assimilation rate were divided by body mass to give mass-specific rates of μWatts/hr/mg worker and mg food/day/mg worker respectively.

#### Statistical Analysis

*Multi-response phylogenetic mixed models*
We used Bayesian multi-response phylogenetic mixed models (MR-PMM) to decompose correlations between species economic traits into phylogenetic and non-phylogenetic components (*20*). Components associated with phylogeny can be thought of as conservative trait correlation (CTC), where-as non-phylogenetic components can be thought of as trait correlations independent of phylogeny *sensu* Westoby, Yates, Holland and Halliwell (*21*). Multi-response models provide a more biologically appropriate model structure than traditional methods (i.e. PGLS, PICs) given that evolutionary selection on traits is often reciprocal rather than unidirectional, and adaptation often proceeds by phylogenetic niche conservatism (*21*). We used trait data collected above from 305 colonies of 123 species from 11 environmental sites. Specifically, we fitted all economic traits jointly as response variables, removing the global intercept to estimate separate intercepts for each response, and using correlated random effects to specify phylogenetic and non-phylogenetic covariance matrices. We modelled multi-level structure in the data by fitting random intercepts across species (accounting for within-species replication), and sites (accounting for the sampling hierarchy). Parameter estimates from models are reported as posterior means with 50% and 95% credible intervals in Fig. 2a and fig. S3. Estimates for which the 95% credible interval did not cross zero are shown as bold lines, while those for which the 70-90% did not cross zero are shown as faded lines, in Fig. 2b and indicated in table S2 and Data S2. Credible intervals represent the likelihood that an effect size is above or below zero and indicate strength of certainty in the relationship. Phylogenetic correlations are estimated with greater uncertainty than standard between-species correlations (*20*), meaning 95% CI are likely to be conservative.

In addition to estimating and decomposing correlations between response variables, we included site-level climate and soil P variables as fixed effect predictors of all response variables except $A_{mass}$ and colony size (derived from collections at a single site and therefore invariant with respect to fixed effect predictors). These were downscaled microclimate mean annual temperature ($MAT_m$) and microclimate mean annual VPD ($VPD_m$) which were interpolated at the site level (11 sites across six locations – see above "Climate data") and soil phosphorus (soil phosphorus) which was measured at the plot level (n = 38 plots, note 3 plots did not have sufficient ants collected for trait measurements). Climate and soil P variables were not significantly correlated (pairwise correlations all $p > 0.05$) and therefore did not pose issues of multicollinearity. All MR-PMM were fit using the MCMCglmm R package (*22*).

A genus-level phylogenetic tree was used to derive the phylogenetic correlation matrix for all analyses. This approach treats species as replicates of a genus when estimating phylogenetic random effects and therefore ignores any phylogenetic structure between species within ant genera. We chose this approach because species level phylogenetic relationships are not well resolved for most Australasian ant taxa. We constructed a genus-level phylogenetic tree using a time-calibrated phylogeny, which includes several representative species of each genus worldwide (*23*). We pruned the tree to the 34 genera included in our study using 'drop.tip' from *ape* ver 5.3 (*24*), and inserted three additional genera using 'bind.tip' from *phytools* (*25*). Genera *Lioponera* and *Zasphinctus* (the only representative genera of Dorylinae included in the study) were placed as sister genera to *Cerapachys* (*26*) (*Cerapachys* was then dropped from the tree). Genus *Chelaner* was placed as sister to *Monomorium* (*27*).

*Model Fitting*

We used parameter expanded priors with (variance) V = $I_k$ (an identity matrix of dimension equal to the number of response traits, k), and (degree of belief) ν = k+1 for random effects (*28*), and default independent normal priors with (mean) = 0 and V = $10^{10}$ for fixed effects. Estimates for all parameters converged successfully with nitts = 110000, burn-in = 10000, and thin = 100. We assessed model convergence from 4 separate MCMC chains by 1) visually inspecting traces of the MCMC posterior estimates; and 2) confirming potential scale reduction factors ($\hat{r}$), a convergence diagnostic test that compares within- and between-chain variance (*29*), were <1.01 for all parameter estimates (fig. S9).

*Model Validation*

We performed model validation using posterior predictive checks and a leave-one-out (LOO) cross-validation (CV) procedure. Specifically, we performed LOO-CV at the species level, by leaving one species out per model fit when calculating the log predictive density. Posterior predictive checks confirm that the fitted model generates plausible data for all response variables (fig. S4). Predictions from LOO-CV show that observed data for left-out species have good coverage at the 95% credible interval and effectively estimate the rank order of species mean phenotypic values (fig. S5). Further, estimates of phylogenetic correlations derived from combining posterior samples across LOO-CV fits had almost identical means and CIs (fig. S5) to those fit to the full dataset (Fig. 2a), indicating that parameter estimates are robust to cross validation. Finally, we explored fitting the model with and without phylogenetic components of trait (co)variance to assess the importance of phylogenetic effects on model predictive performance. A model including phylogenetic (co)variances substantially outperformed a model with no phylogenetic component based on leave one out cross-validation (fig. S10). Furthermore, LOO-CV predictions from this reduced model showed wide CIs and were unable to capture the rank order of species phenotypic values (fig. S11).

*Phylogenetic signal*

We calculated phylogenetic signal in each economic trait from the fitted MR-PMM as λ,

$$\lambda_i = \frac{\sigma_{ii}^{phy}}{\sigma_{ii}^{phy} + \sigma_{ii}^{ind} + \sigma_{ii}^{site} + \sigma_{ii}^{res}}$$

where $\sigma_{ii}^{phy}, \sigma_{ii}^{ind}, \sigma_{ii}^{site}, \sigma_{ii}^{res}$ and $\sigma_{ii}^{res}$ are the estimated phylogenetic, non-phylogenetic, site, and residual variances for trait *i*, respectively.

*Phylogenetic imputation*

Traits varied in the extent of missing data (table S1). Missing response values are permitted in MCMCglmm, with missing values imputed conditional on the full covariance structure of the model. In our case, this means that both phylogenetic and non-phylogenetic trait correlations inform the imputation of missing values (*20, 30*). One benefit of this approach over multiple imputation procedures is that imputation uncertainty is naturally propagated through to the posterior distribution of parameter estimates from the fitted model. We used the gap-filled dataset predicted from the MR-PMM fit to conduct a PCA (see section below and Fig. 3).

*Principle components analysis*

To examine how the economic traits map onto reduced dimensional space, and the spread of major subfamily clades across these dimensions we used principle components analysis. To account for missing data for some traits we used the gap-filled dataset predicted from the MR-PMM fit to calculate species-means for each trait, n = 123 species (full dataset). To examine if the same patterns were present using raw values rather than gap-filled predicted data, we also constructed two datasets using species raw trait values for complete cases of four traits (nitrogen, mass, mass density, mass-specific metabolic rate), n = 65 species (four-traits dataset), and five traits (nitrogen, mass, mass density, mass-specific metabolic rate, lifespan), n = 15 species (five traits dataset). For each of these three datasets we used species trait averages to construct a principle components analysis (PCA) with varimax rotated axes (to aid interpretation) using the function *principle* in the package 'psych' (*31*). We coloured species points by subfamily mapped onto the first two PC axes to observe where the seven major subfamily clades were positioned within multi-trait space. The datasets with four and five traits respectively (that used complete traits without gap-filling data) showed similar axes and overall groupings of subfamilies for these subsets of traits, providing increased confidence that the economic spectrum occurs over two axes of variation (i.e., not an artefact of imputation from MR-PMM model predictions (fig. S12)). We then tested whether variation along the axes of economic strategy was associated with climate and soil P, using linear regressions between PC components and each climate ($T_m$ and $VPD_m$) and soil P variable (table S4, fig. S8).

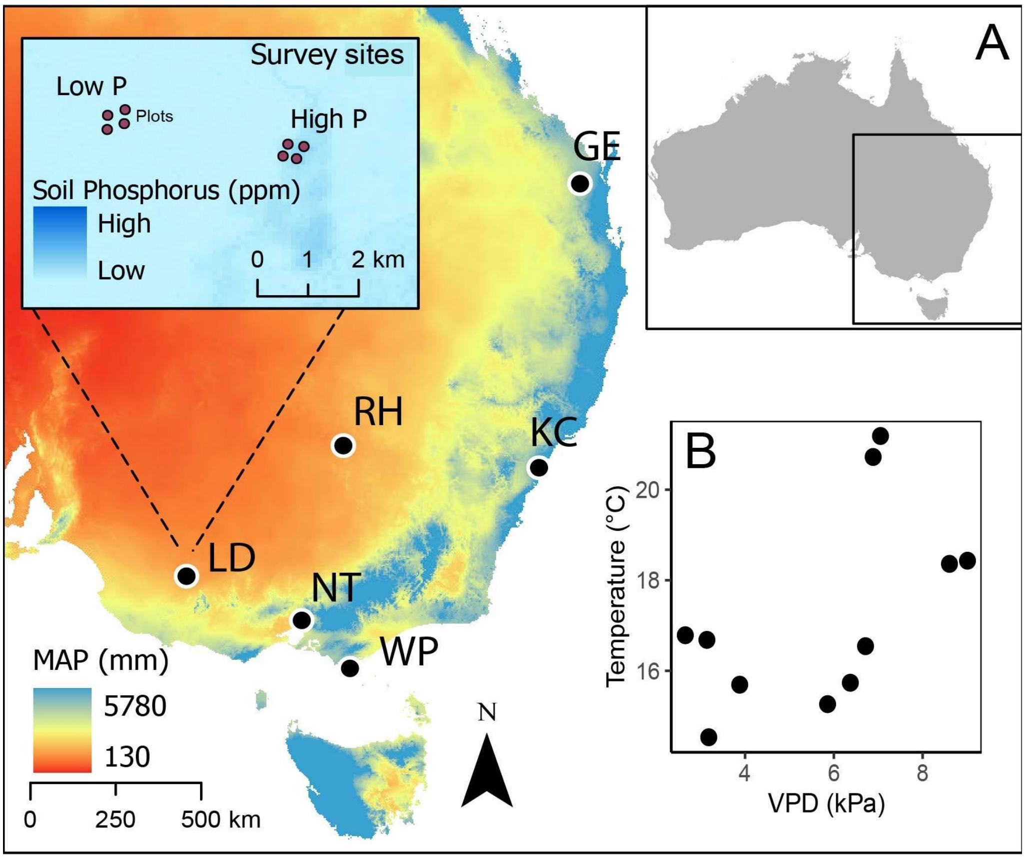


**Fig. S1: Survey design.** A) Map of six locations representing low to high precipitation and temperature across the south-eastern region of Australia, inset indicates layout of sites (high phosphorus and low phosphorus) and plot survey design within each location, all locations were represented by two sites expect for NT represented by one high phosphorus site (total sites = 11), WP = Wilson's Promontory, KC = Ku-ring-gai Chase National Park, GE = Glen Echo, NT = Nangak Tamboree Wildlife Sanctuary, LD = Little Desert National Park, RH = Round Hill Nature Reserve, B) Mean annual microclimate temperature ($T_m$) and microclimate vapor pressure deficit ($VPD_m$), representing aridity for eleven sites.

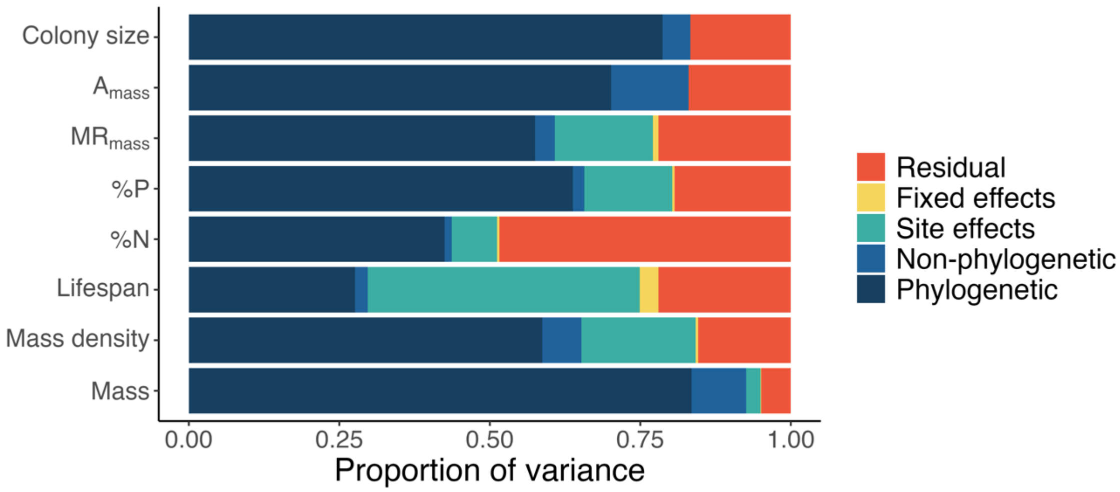


**Fig. S2: Decomposition of trait variance (partial $R^2$) attributed to different levels in the model hierarchy.** Phylogenetic = phylogenetically structured between-genus effects. Non-phylogenetic = non-phylogenetic (independent) between-species effects. Site effects = between survey locations. Fixed effects = variables of site-specific mean annual microclimate temperature, mean annual microclimate vapor pressure deficit (representing aridity), and soil phosphorus. Residual = residual variance unexplained by the model.

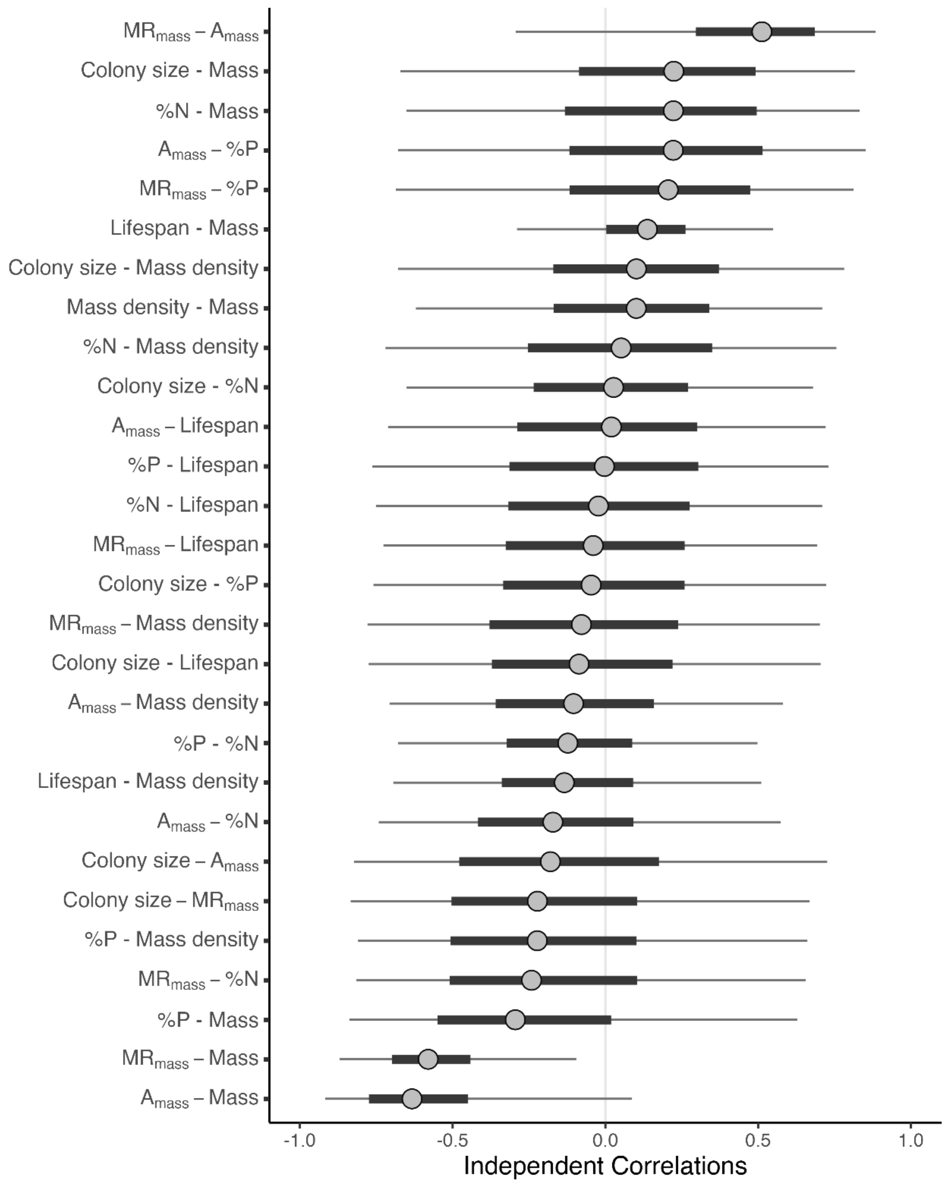


**Fig. S3. Posterior summaries of independent (non-phylogenetic) correlation coefficients from multi-response phylogenetic mixed model.** Showing the posterior mean (point), 50% (heavy wick) and 95% (light wick) credible intervals.

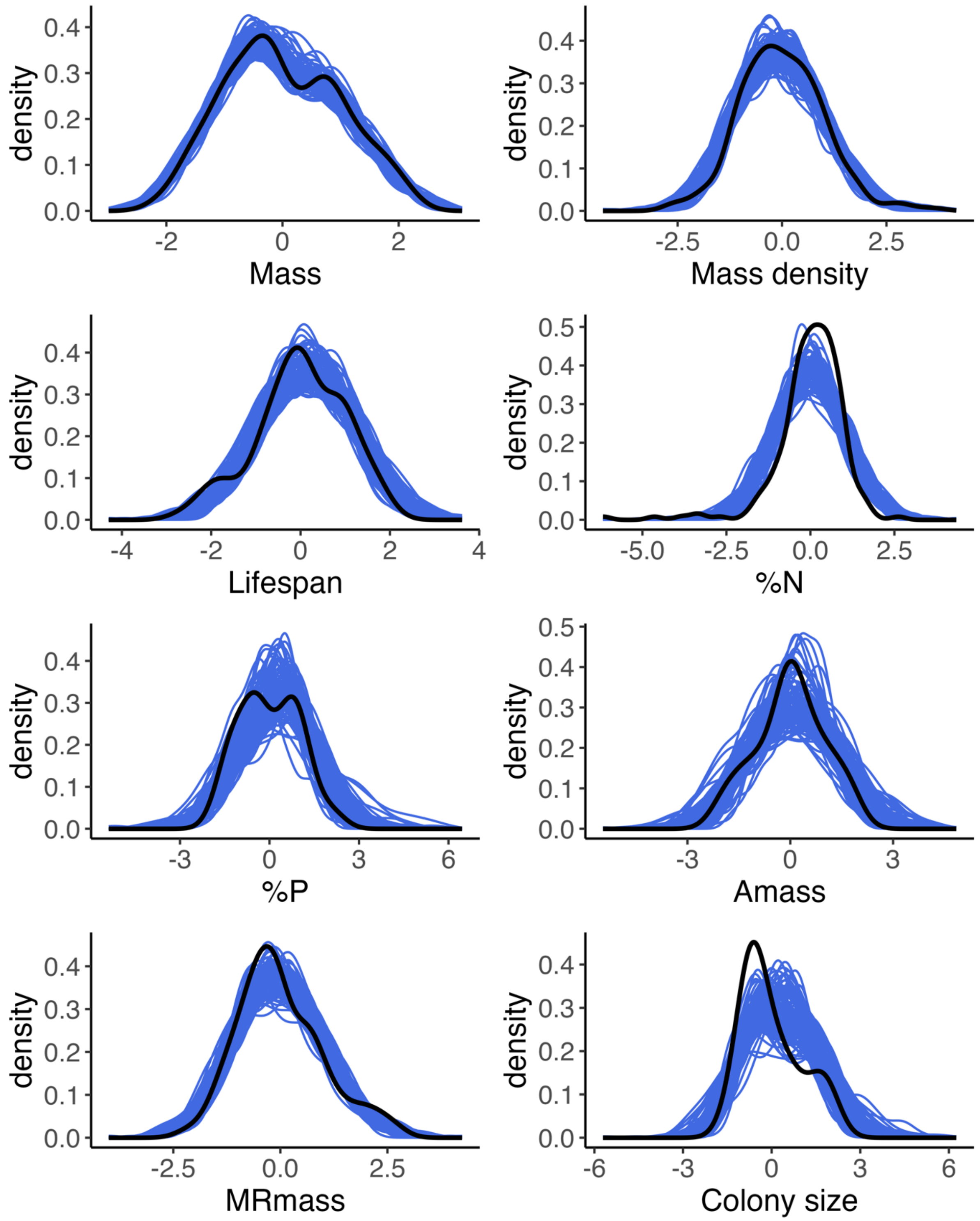


**Fig. S4: Posterior predictive checks.** Posterior predictive checks (draws = 100) from the MR-PMM fit show good alignment between predicted data (in blue) and raw data (in black).

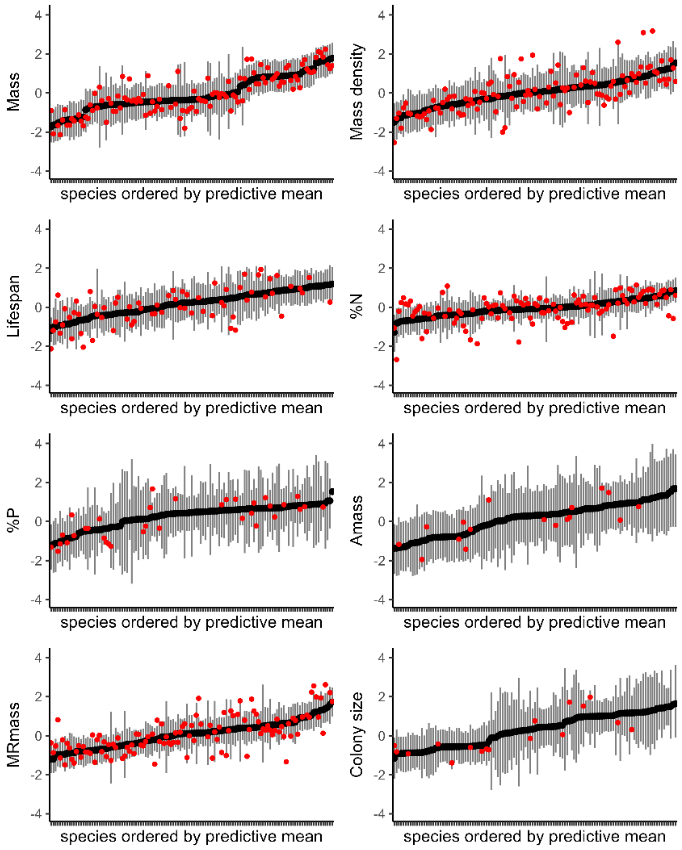


**Fig. S5: Observed species mean trait values (red points) superimposed over model predictions for left-out species based on LOO-CV.** Black points indicate the posterior predictive mean for each species, grey bars indicate the 95% credible intervals. X-axes are ordered by the predictive mean to assess the capacity of the models to predict the rank order of observed values. Predictions from the main model including phylogenetic components effectively capture the rank order of observations.

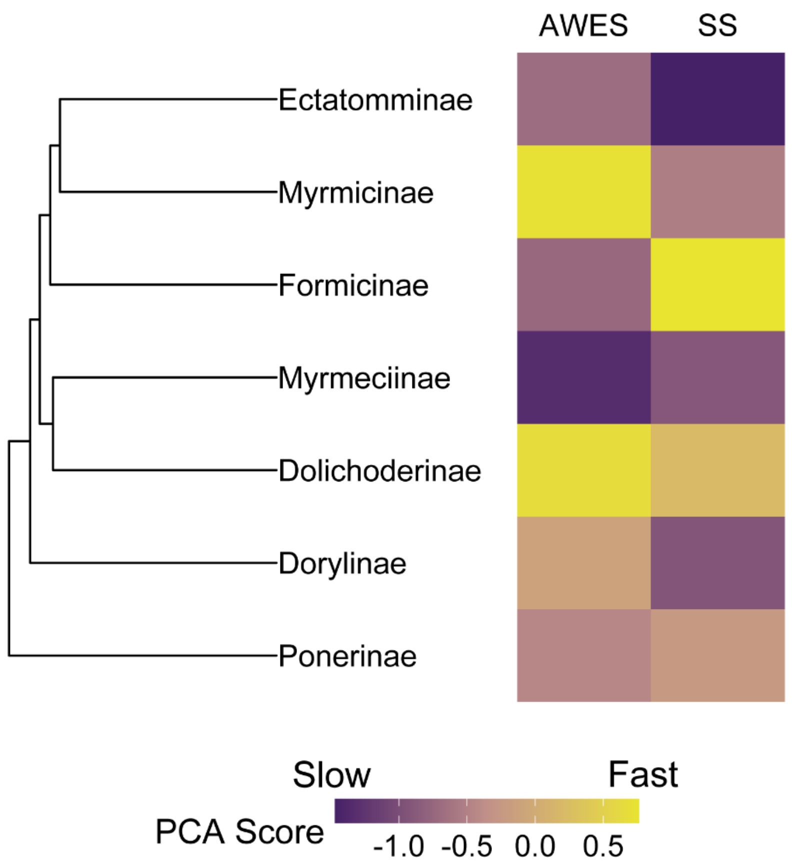


**Fig. S6: Mapping the superorganism economic spectrum onto the ant phylogenetic tree at the subfamily level.** Showing PCA scores from Fig. 3. For axis 1: Ant Worker Economic Spectrum (AWES) and for axis 2: Stoichiometry Spectrum (SS). Slow values (darker colours) for AWES signify large body mass, long lifespan, high mass density, low mass-specific metabolic and assimilation rates and small colony sizes. Fast values (lighter colours) represent the opposite values for each trait. Slow values (darker colours) for stoichiometry spectrum (SS) represent high nitrogen concentration and low phosphorus concentration, with the opposite ratio for fast ecological strategies (lighter colours). This demonstrates that niche partitioning is occurring along two axes of the ant economic spectrum (see PCA Fig. 3 to see these patterns mapped onto multidimensional space).

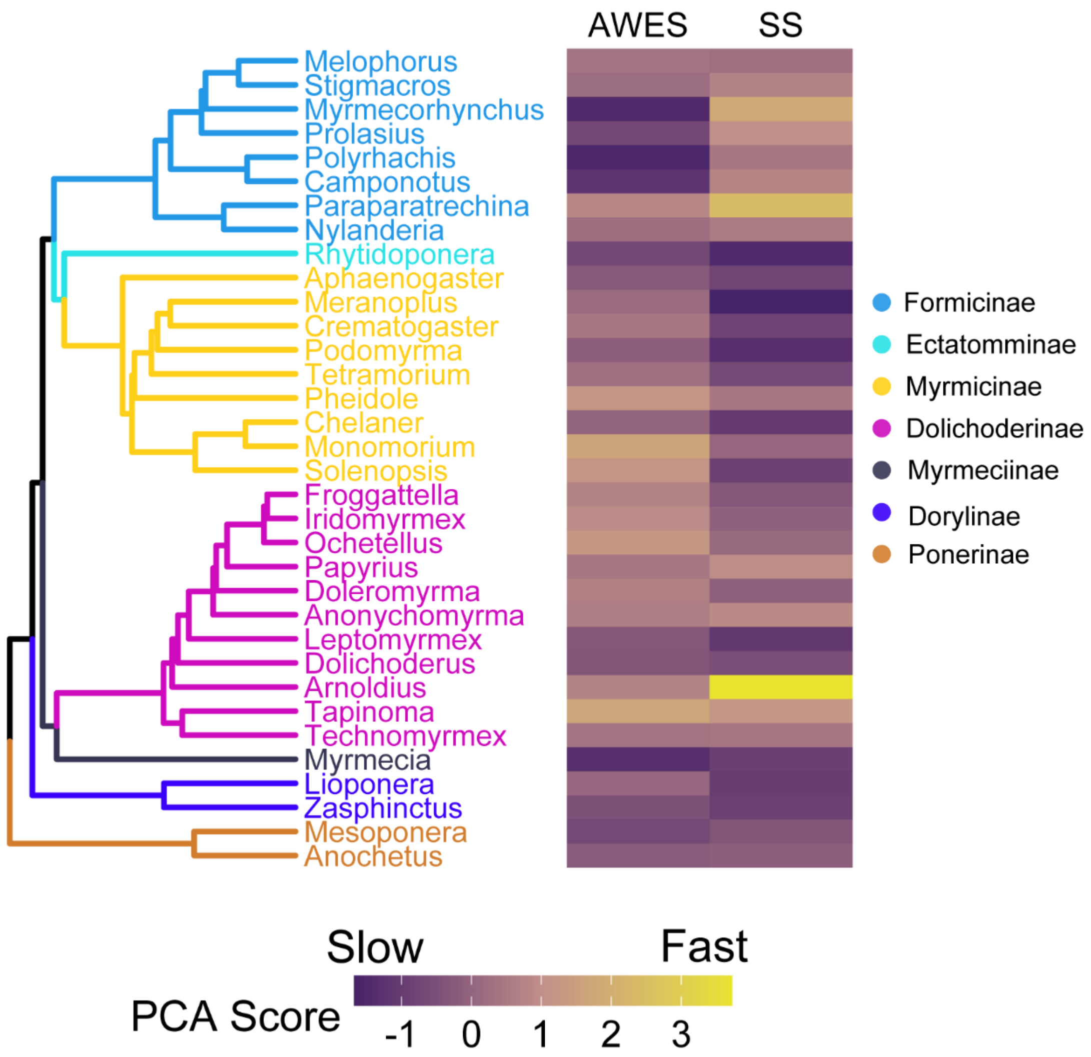


**Fig. S7: Mapping the superorganism economic spectrum onto the ant phylogenetic tree at the genus level including genus *Arnoldius*.** Showing PCA scores from Fig. 3 main text. For axis 1: Ant Worker Economic Spectrum (AWES) and for axis 2: Stoichiometry Spectrum (SS). Slow values (darker colours) for AWES signify, large body mass, long lifespan, high mass density, low mass-specific metabolic and assimilation rates and small colony sizes. Fast values (lighter colours) represent the opposite values for each trait. Slow values (darker colours) for stoichiometry spectrum (SS) represent high nitrogen concentration and low phosphorus concentration, with the opposite ratio for fast ecological strategies (lighter colours). This demonstrates that niche partitioning is occurring along two axes of the superorganism economic spectrum (see PCA Fig. 3 to see this mapped onto multidimensional space).

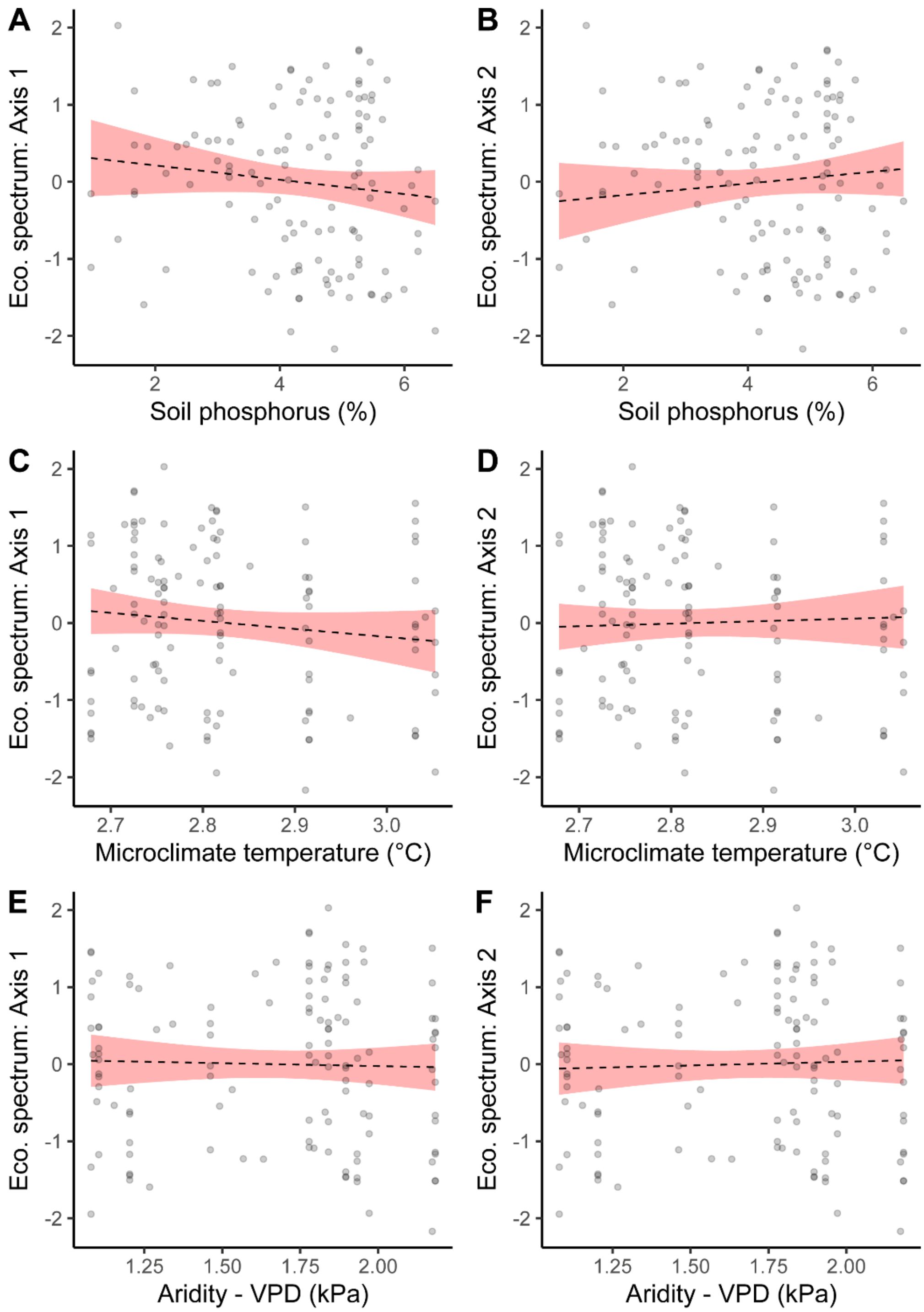


**Fig. S8: The worker economic spectrum does not vary with climate and soil phosphorus variables tested here.** Showing varimax-rotated PCA axes 1 and 2 representing two dimensions of the worker economic spectrum with climate and soil phosphorus as predictor variables, points represent species, lines and 95% CI from model fits (Table S5). Microclimate temperature (MATm) is mean annual microclimate. Aridity gradient represented by microclimate vapor pressure deficit (VPDm).

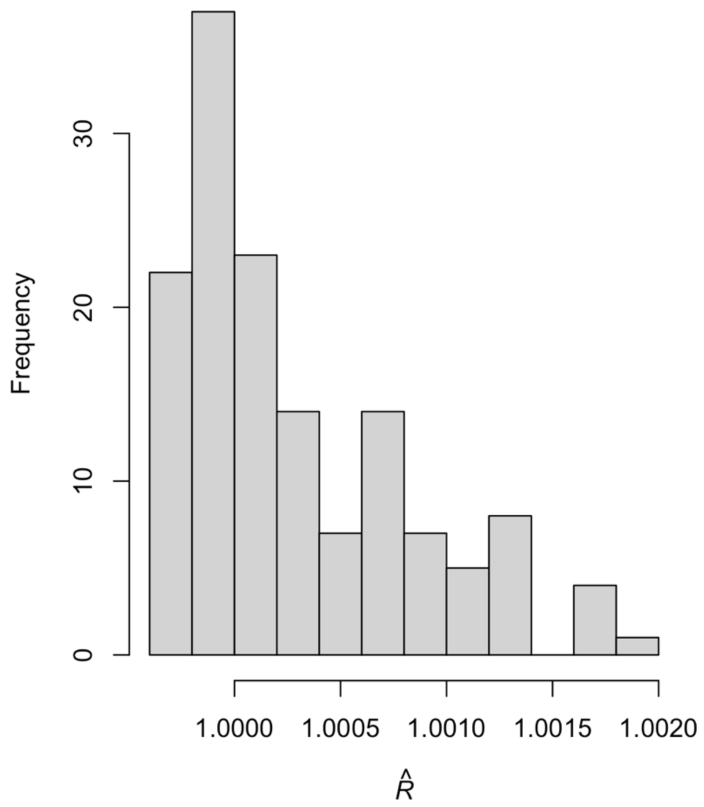


**Fig. S9: $\hat{R}$ values for MR-PMM model.** Shows that $\hat{R}$ values for all variance parameters in the model were <1.01 indicating successful convergence of MCMC chains.

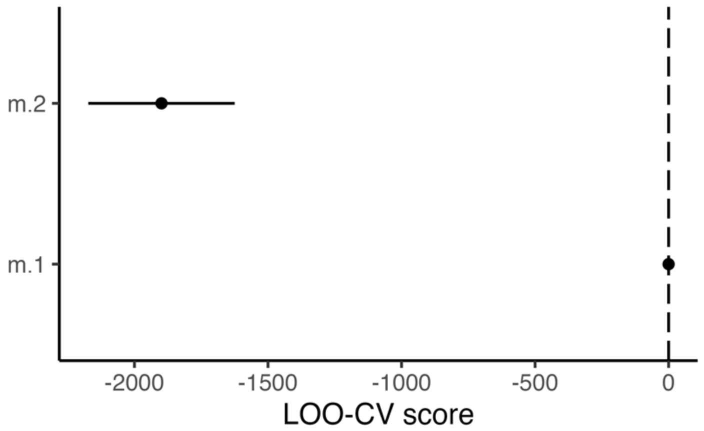


**Fig. S10: Testing a MR-PMM model without including phylogenetic effects.** A model including phylogenetic trait (co)variances (m.1) substantially outperforms a model excluding phylogenetic trait (co)variances (m.2) based on LOO-CV scores.

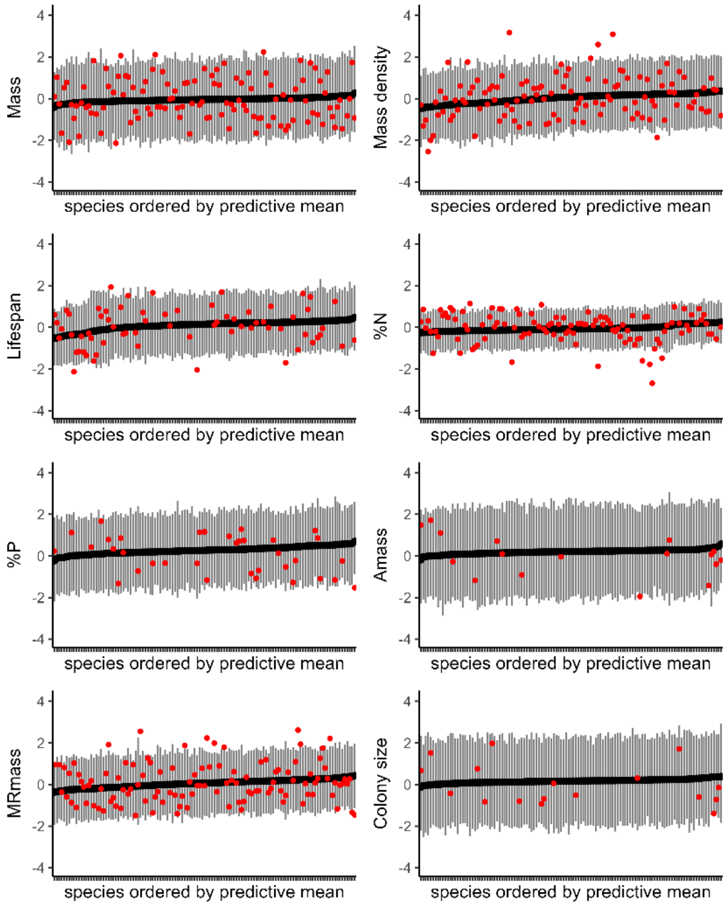


**Fig. S11: Observed species mean trait values (red points) superimposed over model predictions, for a model without including phylogenetic components, for left-out species based on LOO-CV.** Black points indicate the posterior predictive mean for each species, grey bars indicate the 95% credible intervals. X-axes are ordered by the predictive mean to assess the capacity of the models to predict the rank order of observed values. Predictions from the model without phylogenetic components have a higher uncertainty and fail to capture the rank order of observed values compared to the model including phylogenetic components (fig. S5).

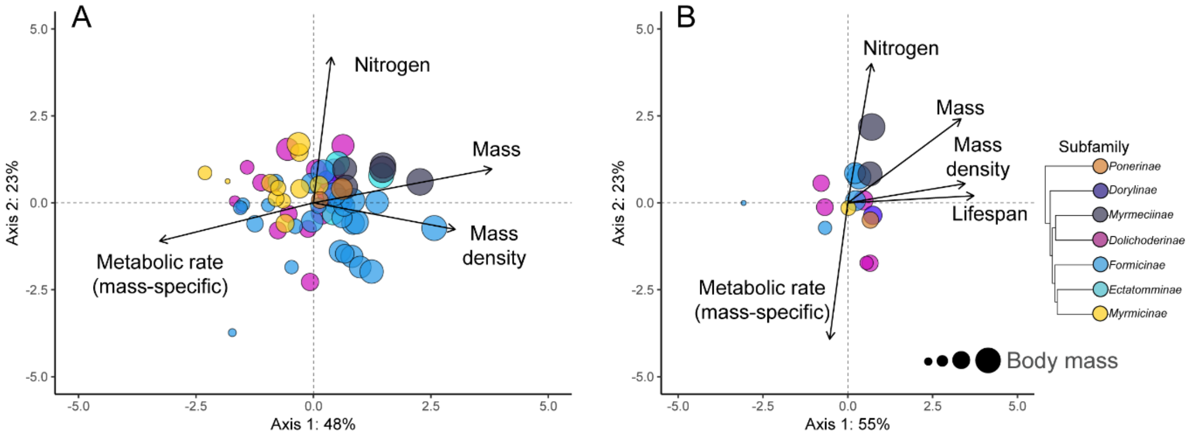


**Fig. S12: Mapping the superorganism economic spectrum onto multidimensional space using subsets of traits with full data coverage.** Showing varimax-rotated PCA from A) raw species-averaged trait data for four traits representing complete cases (no missing data) from 65 species and B) raw species-averaged trait data for five traits representing complete cases (no missing data) from 15 species coloured as subfamily groups. Percentage variation explained in brackets for axis 1 and axis 2. Size of points represents dry body mass scaled between 1-10. Ant subfamily phylogenetic tree with subfamily colours as legend.

**Table S1: Economic traits of worker ants**. Showing abbreviations (term), units, % trait coverage across species sampled (n = 123), and equivalent trait in the Leaf Economics Spectrum (LES).

| Trait | Term | Units | % coverage | Parallel to LES |
|---|---|---|---|---|
| Worker mass density | Mass density | $mg/mm^3$ | 94.31 | Leaf mass per unit area |
| Mass-specific assimilation rate | $A_{mass}$ | mg/day/mg | 13.82 | Photosynthetic capacity |
| Worker N concentration | N | % | 92.68 | Leaf N concentration |
| Worker P concentration | P | % | 28.46 | Leaf P concentration |
| Mass-specific metabolic rate | $MR_{mass}$ | µW/mg | 91.06 | Dark respiration |
| Lifespan | Lifespan | days | 54.47 | Lifespan |
| Colony size | Colony size | No. workers | 13.82 | NA |
| Body mass | Mass | mg | 94.31 | NA |

**Table S2. Mean posterior correlation coefficients from multi-response phylogenetic mixed model.** Showing the phylogenetic correlation (upper triangular) and independent correlation (lower triangular) of each pairwise trait combination. Significance at 95% credible interval (CI) indicated by *, correlations significant at 70-90% CI shown in bold.

| | Mass | Mass density | Lifespan | % N | % P | $A_{mass}$ | $MR_{mass}$ | Colony size |
|---|---|---|---|---|---|---|---|---|
| Mass | | **0.60*** | **0.57*** | **0.68*** | **-0.46** | **-0.72*** | **-0.86*** | **-0.70*** |
| Mass density | 0.13 | | **0.42** | **0.49*** | **-0.41** | **-0.54** | **-0.63*** | **-0.49** |
| Lifespan | 0.09 | -0.09 | | **0.55*** | **-0.43** | **-0.49** | **-0.61*** | **-0.63*** |
| % N | 0.16 | 0.02 | 0.01 | | **-0.57** | **-0.51** | **-0.76*** | **-0.67*** |
| % P | -0.24 | -0.15 | -0.02 | -0.07 | | **0.35** | **0.54** | **0.51** |
| $A_{mass}$ | **-0.58** | -0.12 | -0.009 | -0.14 | 0.18 | | **0.67** | **0.57** |
| $MR_{mass}$ | **-0.55*** | -0.11 | -0.04 | -0.19 | 0.17 | **0.46** | | **0.74*** |
| Colony size | 0.18 | 0.08 | -0.03 | 0.04 | -0.07 | -0.18 | | |

**Table S3: Phylogenetic signal of economic spectrum traits.** Posterior quantiles (0.5, 0.025, 0.975) of phylogenetic signal (λ) for each economic trait from the MR-PMM model.

| Trait | λ (±95% CI) |
|---|---|
| Mass | 0.84 (0.69 – 0.91) |
| Mass density | 0.63 (0.29 – 0.80) |
| Lifespan | 0.36 (0.09 – 0.63) |
| Phosphorus | 0.65 (0.23 – 0.89) |
| Nitrogen | 0.41 (0.16 – 0.66) |
| $A_{mass}$ | 0.68 (0.30 – 0.89) |
| $MR_{mass}$ | 0.62 (0.28 – 0.78) |
| Colony size | 0.77 (0.42 – 0.93) |

**Table S4: Testing the effect of environmental variables on the worker economic spectrum.** Linear OLS models of varimax-rotated PCA axes 1 and 2 representing two dimensions of the worker economic spectrum with climate and soil phosphorus as predictor variables.

| Model | Adj. $R^2$ | Coef. (± SE) | p-value |
|---|---|---|---|
| Axis 1 ~ $\log_{10}$ (Soil P) | 0.01 | -0.10 (0.07) | 0.153 |
| Axis 2 ~ $\log_{10}$ (Soil P) | 0.01 | 0.09 (0.07) | 0.192 |
| Axis 1 ~ $MAT_m$ | 0.01 | -1.05 (0.81) | 0.199 |
| Axis 2 ~ $MAT_m$ | 0.00 | 0.23 (0.82) | 0.783 |
| Axis 1 ~ $VPD_m$ | 0.00 | -0.09 (0.24) | 0.707 |
| Axis 2 ~ $VPD_m$ | 0.00 | 0.12 (0.24) | 0.611 |

**Data S1. (separate file)**

**Model output for MR-PMM model.** Phylogenetic and non-phylogenetic trait correlations showing the posterior predictions quantiles (0.5 (mean), 0.025, 0.975) from the MR-PMM model and the significance of the relationships based on credible intervals at 80% CI, 85% CI, 90% CI, and 95% CI, see also Fig. 2a for visualization of credible intervals.

**Data S2. (separate file)**

**Model summary for MR-PMM model.** The model summary for each component in the MR-PMM model including fixed effects (environmental variables). See code and data repository for further information regarding model construction.

#### References


1. R. Viscarra Rossel *et al.*, "Soil and Landscape Grid National Soil Attribute Maps - Total Phosphorus (3" resolution) - Release 1. v6.," *Data Collection* (CSIRO, 2014).
2. G. E. Rayment, D. J. Lyons, *Soil chemical methods: Australasia*. (CSIRO publishing, 2011), vol. 3.
3. M. R. Kearney, P. K. Gillingham, I. Bramer, J. P. Duffy, I. M. D. Maclean, A method for computing hourly, historical, terrain-corrected microclimate anywhere on earth. *Methods in Ecology and Evolution* **11**, 38-43 (2020).
4. I. M. D. Maclean, J. R. Mosedale, J. J. Bennie, Microclima: An r package for modelling meso- and microclimate. *Methods in Ecology and Evolution* **10**, 280-290 (2019).
5. M. U. Kemp, E. Emiel van Loon, J. Shamoun-Baranes, W. Bouten, RNCEP: global weather and climate data at your fingertips. *Methods in Ecology and Evolution* **3**, 65-70 (2012).
6. J. Hollister, T. Shah, M. Beck, Zenodo, Ed. (2017).
7. J. Monteith, M. Unsworth, *Principles of environmental physics: plants, animals, and the atmosphere*. (Academic press, 2013).
8. E. Csata, A. Dussutour, Nutrient regulation in ants (Hymenoptera: Formicidae): a review. *Myrmecological News* **29**, 111-124 (2019).
9. J. R. Lighton, *Measuring metabolic rates: a manual for scientists*. (Oxford University Press, 2018).
10. D. Bates, M. Maechler, B. M. Bolker, S. Walker, Fitting Linear Mixed-Effects Models Using lme4. *Journal of Statistical Software* **67**, 1-48 (2015).
11. S. L. Chown *et al.*, Discontinuous gas exchange in insects: a clarification of hypotheses and approaches. *Physiological and Biochemical Zoology* **79**, 333-343 (2006).
12. T. Therneau, A Package for Survival Analysis in R. **R package version 3.5-5**, (2023).
13. C. L. Parr *et al.*, GlobalAnts: a new database on the geography of ant traits (Hymenoptera: Formicidae). *Insect Conservation and Diversity* **10**, 5-20 (2017).
14. California Academy of Science. (California Academy of Science, 2024), vol. 2024.
15. J. Schindelin *et al.*, Fiji: an open-source platform for biological-image analysis. *Nature Methods* **9**, 676-682 (2012).
16. N. Blüthgen, G. Gebauer, K. Fiedler, Disentangling a rainforest food web using stable isotopes: dietary diversity in a species-rich ant community. *Oecologia* **137**, 426-435 (2003).

17. J. T. Buxton, K. A. Robert, A. T. Marshall, T. L. Dutka, H. Gibb, A cross-species test of the function of cuticular traits in ants (Hymenoptera: Formicidae). *Myrmecological News* **31**, 31-46 (2021).
18. D. G. Chapman, Some properties of the hypergeometric distribution with applications to zoological censuses. *Univ. Calif. Stat.* **1**, 60-131 (1951).
19. B. A. Krabbe *et al.*, Using nutritional geometry to define the fundamental macronutrient niche of the widespread invasive ant Monomorium pharaonis. *PLoS One* **14**, e0218764 (2019).
20. B. Halliwell, B. R. Holland, L. A. Yates, Multi-response phylogenetic mixed models: concepts and application. *Biological Reviews* **100**, 1294-1316 (2025).
21. M. Westoby, L. Yates, B. Holland, B. Halliwell, Phylogenetically conservative trait correlation: Quantification and interpretation. *Journal of Ecology* **111**, 2105-2117 (2023).
22. J. D. Hadfield, MCMC methods for multi-response generalized linear mixed models: the MCMCglmm R package. *Journal of statistical software* **33**, 1-22 (2010).
23. E. P. Economo, N. Narula, N. R. Friedman, M. D. Weiser, B. Guénard, Macroecology and macroevolution of the latitudinal diversity gradient in ants. *Nature Communications* **9**, 1778 (2018).
24. E. Paradis, K. Schliep, ape 5.0: an environment for modern phylogenetics and evolutionary analyses in R. *Bioinformatics* **35**, 526-528 (2019).
25. L. J. Revell, phytools: an R package for phylogenetic comparative biology (and other things). *Methods in ecology and evolution* **3**, 217-223 (2012).
26. M. L. Borowiec, Generic revision of the ant subfamily Dorylinae (Hymenoptera, Formicidae). *ZooKeys*, 1 (2016).
27. K. S. Sparks, A. N. Andersen, A. D. Austin, A multi-gene phylogeny of Australian *Monomorium* Mayr (Hymenoptera : Formicidae) results in reinterpretation of the genus and resurrection of *Chelaner* Emery. *Invertebrate Systematics* **33**, 225-236 (2019).
28. P. de Villemereuil, Quantitative genetic methods depending on the nature of the phenotypic trait. *Annals of the New York Academy of Sciences* **1422**, 29-47 (2018).
29. A. Vehtari, A. Gelman, D. Simpson, B. Carpenter, P.-C. Bürkner, Rank-normalization, folding, and localization: An improved Rˆfor assessing convergence of MCMC (with discussion). *Bayesian analysis* **16**, 667-718 (2021).
30. P. Sanchez-Martinez *et al.*, A framework to study and predict functional trait syndromes using phylogenetic and environmental data. *Methods in Ecology and Evolution* **15**, 666-681 (2024).
31. W. Revelle, psych: Procedures for Psychological, Psychometric, and Personality Research. *https://CRAN.R-project.org/package=psych*, (2024).
